# Divergent morphologies with convergent performance in the mandible of pelagiarian fishes

**DOI:** 10.1101/2025.01.28.634496

**Authors:** Andrew Knapp, Gizeh Rangel-de Lazaro, Matt Friedman, Zerina Johanson, Kory M Evans, Sam Giles, Hermione T Beckett, Anjali Goswami

## Abstract

Mandibles represent a key evolutionary innovation that enabled jawed vertebrates to adapt and diversify in response to a range of food sources. Using a phylogenetic comparative approach, we explore the phenotypic disparity and mechanical properties of the lower jaw in Pelagiaria, a morphologically diverse but relatively small clade of open-ocean fishes which are hypothesized to have radiated near the Cretaceous/Paleogene (K/Pg) mass extinction event. We found that body elongation and diet are not significantly correlated with jaw shape, but that habitat depth and tooth type are. Mechanical advantage (MA) is significantly correlated with mandible shape, with jaw-closing MA being most strongly correlated. Pelagiarian jaw shapes fall broadly into six morphotypes, of which two show significantly higher closing MA than other groups, despite differing substantially in shape. The high morphological disparity of pelagiarian mandible shape was established very early in their evolutionary history, and high levels of disparity have been maintained over tens of millions of years; this is consistent with the hypothesis that Pelagiaria represents an ancient adaptive radiation. Our results demonstrate both the mechanical and morphological diversity of the pelagiarian mandible and highlight the crucial role that morphological diversification has played in the trophic radiation of this clade.

## Introduction

The lower jaw (mandible) is an indisputably critical component of the vertebrate skull, with a primary role in acquiring and processing food. The evolution of jaws during or before the early Silurian (Andreev et al., 2022; Zhu et al. 2022) is considered a key innovation in vertebrate evolution (DeLaurier and Gerhart, 2019), with more than 99% of the living ∼70 000 species of backboned animals belonging to Gnathostomata, the jawed vertebrates. Gnathostomes have evolved to occupy a wide range of ecological roles in both terrestrial and aquatic environments worldwide, and the form of the lower jaw has consequently adapted to reflect the diverse feeding strategies associated with these disparate ecologies (Coombs et al., 2024; Deakin et al., 2022; Hill et al., 2018; Morales-Garcia et al., 2021). Jaw morphology is closely linked with feeding ecology due to physical constraints that set limits on performance variables such as bite force, jaw velocity, and, in aquatic taxa, suction flow (Westneat, 2004). Evolutionary radiations that are characterised by trophic diversifications are therefore expected to be accompanied by diversification of feeding strategies, which should in turn be reflected in jaw shape.

Teleosts account for around 96% of all extant fishes and roughly half of extant vertebrates (Nelson et al., 2016). Their evolutionary success and global distribution have been attributed to their ability to adapt to every marine and freshwater environment, and they have evolved to exploit a wide range of diets (Pavlov and Kasumyan, 2002). The teleost mandible is made up of three elements: the tooth-bearing dentary, and the postdentary angular and articular (sometimes fused as single anguloarticular), which form the jaw joint with the quadrate and act as the main anchor point for the muscles and tendons involved in mandibular movement (Datovo and Vari, 2013; Gregory, 1932; Winterbottom, 1973). The elements of the oral jaw articulate with the suspensorium and neurocranium to make up the skull and, in many teleost taxa, form part of a kinetic four-bar linkage system in conjunction with a highly mobile upper jaw composed of the maxilla and premaxilla (Motta, 1984; Westneat, 1990). This structurally complex apparatus has evolved to enable suction feeding via buccal expansion to capture prey in the relatively dense and viscous aquatic environment (Day et al., 2015; Ferry-Graham and Lauder, 2001, Westneat, 2004). Teleost jaws contrast with those of tetrapods, in which the mandible is usually the only kinetic component, and the upper jaw is generally akinetic and fused with the cranium, although notable examples of limited cranial kinesis are observed in birds (Zusi, 1984) and squamates (Frazzeta, 1962) and in the intercranial hinge of some lagomorphs (Wood-Bailey et al., 2022).

Despite the obvious complexities of the teleost feeding mechanism, past work shows that lower jaw function can be examined through simple biomechanical models. The teleost mandible acts as a first-class lever when opening and a third-class lever when closing. Effort is exerted between the fulcrum (quadrato-mandibular joint) and the in-lever, either the attachment point of the adductor mandibularis on the coronoid process of the angular for closing, or the attachment point of the preopercular tendon on the posterior tip of the angular (sometimes a separate retroarticular ossification or process) for opening (Albertson et al., 2005; Westneat, 2004). Mechanical advantage (MA) is a measure of the force amplification achieved through a lever (i.e., the mandible), calculated as the ratio of in-lever length to out-lever length (MA = L_in_ /L_out_). Although a simplified model of jaw closing, it provides a useful comparative functional variable and is commonly used to assess biting performance across teleosts and other vertebrates (Albertson et al., 2005; Alfaro et al., 2004; Anderson et al., 2011; Burress et al., 2020; Deakin et al., 2022; Hampton, 2011; Heiple et al., 2023; Morales-Garcia et al., 2021; Singh et al., 2021; Wainwright et al., 2004; Westneat, 1994, 1995, and 2004). Higher MA values imply higher force transmission, with a trade-off against angular velocity (i.e., movement speed), although a 2012 study found no relationship between jaw opening MA and speed of jaw depression during opening in serranid fishes (Oufiero et al., 2012). The connected elements of the oral jaws have been found to be highly integrated in several clades (Conith et al., 2020; Conith and Albertson, 2021; Jamniczky et al., 2014; Larouche et al., 2022). In some teleost taxa the lower jaw is linked to the maxilla via the coronoid process, where it forms the kinetic input of a four-bar linkage system that comprises the mandible, maxilla, suspensorium and neurocranium (Motta, 1984; Westneat, 1990), causing the premaxilla to rotate anteriorly when the lower jaw is depressed. In these taxa, the output of force transmission when opening the oral jaws may therefore primarily be via the coronoid process.

Mandibular proportions are thus intimately connected to mechanical performance, with elongate, slender jaws tending to have lower MA (low force transmission, high velocity) and short, deep jaws tending to have higher MA (high force transmission, low velocity; Albertson et al., 2005). The mode of feeding in teleosts is consequently expected to reflect this trade-off; ambush predators and suction feeders require high mandible velocities to capture prey and thus lower MA, while taxa that rely on high bite forces are expected to have higher MA (Westneat, 2004). The relationship between mandible length to in-lever length appears straightforward but may have differing functional implications because of the structural and functional complexity of the teleost oral jaws. Shorter mandibles may enable the formation of a planar and near-circular mouth shape, which can improve flow rates into the mouth when suction feeding (Skorczewski et al., 2012). Conversely, taxa that feed on mobile prey often evolve elongate, gracile mandibles for improved suction feeding because more elongate jaws tend to result in a lower MA value and therefore faster velocity transmission (Albertson et al., 2005). Beside mechanical performance, other factors may influence jaw shape. For example, relatively longer jaws allow a larger gape, thus enabling the capture of larger prey.

Mandible shape is diverse and complex in vertebrates, and a single variable such as MA may provide limited information on the drivers of mandible evolution because alternative morphologies can yield equivalent functions (Alfaro et al., 2004). Geometric morphometrics has been used to explore rates of shape evolution, phenotypic disparity, and modularity and integration across a wide range of elements and taxa (Adams et al., 2013; Coombs et al., 2024; Goswami et al., 2019; Knapp et al., 2023). The analysis of shape can therefore provide useful contextual information on the coevolution of form and function when exploring possible adaptive radiations (Deakin et al., 2022). Here, we investigate mandible shape evolution in Pelagiaria, a monophyletic group of open-ocean fishes, to explore the relationships between shape, mechanical performance, and ecology. Paleontological data and time-calibrated phylogenies indicate Pelagiaria underwent rapid diversification around the Cretaceous/Paleogene (K/Pg) mass extinction event (Friedman, 2009 & 2010; Miya et al., 2013), thought to be driven by adaptative radiation to fill recently vacated ecological niches in the open ocean (Friedman et al., 2019). Pelagiaria comprises 286 extant species in 16 families (Friedman et al., 2019; Miya et al., 2013), but, despite its relatively low taxonomic diversity, body shape has been shown to span the continuum of shapes found across teleosts (Friedman et al., 2019; Knapp et al., 2023).

Furthermore, pelagiarian families differ so greatly in morphology that before the advent of molecular phylogenetics they were distributed across as many as six distantly related suborders (Miya et al., 2013; Pastana et al., 2021). Due to the exclusively open ocean habitat of Pelagiaria there are no durophagous or herbivorous taxa, as are commonly found in taxa associated with coral reefs and other shallow water habitats (e.g., Labridae; Corn et al., 2022), and most pelagiarian taxa are opportunistic, being either predators or planktivorous. Despite their relatively narrow dietary breadth, pelagiarian fishes have evolved diverse feeding strategies including fast pursuit predators (e.g., tuna, *Thunnus*), ram suspension feeders (e.g., mackerel, *Scombrus*), deep-sea ambush predators/engulfers (e.g., swallowers, Chisamodontidae), and specialized feeding on gelatinous zooplankton, i.e., salps or the tentacles and gonads of siphonophores (e.g., driftfish, Nomeidae). These feeding strategies are likely to be reflected in jaw shape, which can be a reliable measure of functional diversification in the absence of direct observation, which is not possible for the majority of pelagiarian taxa. If the origin of this clade’s diversity is due to adaptive radiation to fill trophic niches in the open ocean, this may be reflected in the evolution of mandibular shape and mechanical performance, with early and rapid accumulation of morphological and functional disparity. Furthermore, we expect feeding mode, rather than diet, to have a stronger influence on mandible morphology, and we predict that tooth type and jaw shape are likely to be correlated because of the importance of tooth morphology and feeding biology (Mihalitis and Bellwood, 2019).

## Methods

### Data collection, imaging and landmarking

We created high resolution 3D models of the mandible using X-ray micro-computed tomography (μCT) for 137 spirit-preserved pelagiarian species from museum collections (Supplementary Table S1). These specimens represent 15 of 16 pelagiarian families and 62 of 75 genera (Supplementary Table S1 and Supplementary Fig. S2).

Specimens were scanned with pixel size and slice thickness ranging between 0.01 and 0.5 mm and an average of 1657 projections for each specimen. We segmented the lower jaw of each specimen and exported it as a polygon mesh in ASCII PLY format with Materialise Mimics 21.0 (Materialise, 2015), and meshes were decimated to ∼300,000 polygons in Geomagic Wrap 2017 (3D Systems Inc., Rock Hill, SC, USA). We devised a landmark scheme to capture the shape of both the lingual and labial surfaces of the lower jaw, comprising 8 anatomical (Bookstein Type I) landmarks and 10 semi-landmark curves, totalling 223 landmarks per specimen. All landmarks and curves were applied with Stratovan Checkpoint (v. 2020.10.13.0859) on the left hemimandible only (Supplementary Figure S3). We exported landmarks to R Studio (RStudio Team, 2020) and slid curves to minimise bending energy (Zelditch et al., 2004). We aligned landmark data for all specimens with a generalised Procrustes alignment (GPA; Rohlf & Slice, 1990) to minimise scale, position, and rotational differences between specimens, using the ‘gpagen’ function in *geomorph* (Adams et al., 2023). We then performed a principal component analysis (PCA) on the Procrustes-aligned data to explore shape variation within the dataset.

### Phylogeny

We added 23 taxa to the dated phylogeny from Knapp et al. (2023), which was based on Friedman *et al*. (2019). First, we substituted species in our dataset which were not included in Friedman et al. with congeneric species or grafted them to the phylogeny based on other published phylogenies (Arcila et al., 2021; Beckett et al., 2018; Jeena et al., 2022; Miya et al., 2013). We placed new nodes arbitrarily at the midpoint of the bisected branch. Where new taxa were grafted to the stem of established clades, we placed nodes at a distance from the crown node equivalent to the shortest of the crownward branches stemming from that crown node (Supplementary Fig. S1).

### Ecological data

We collected ecological information for each taxon to investigate correlations of mandible shape with ecology. Firstly, we assessed evolutionary allometry by performing a multivariate regression of mandible shape against centroid size using the ‘procd.pgls’ function in *geomorph* (Adams et al., 2013). Body elongation is a dominant form of morphological disparity across teleosts (Claverie and Wainwright, 2014) and is strongly correlated with neurocranium shape in Pelagiaria (Collar et al., 2022; Knapp et al., 2023). We gathered lateral body elongation data for all specimens using the measurement protocol outlined in Claverie and Wainwright (2014; fineness ratio = standard length/body depth). With this method, a circular body has a value of 1 and an elongate body has a value of >>1 (Supplementary Table S1). Following Martinez et al. (2021), we assigned each species to one of three depth zones: *Shallow,* equivalent to the epipelagic zone (0-200m, n = 44); *Intermediate*, equivalent to the mesopelagic zone (200-1000m, n = 51); and *Deep*, equivalent to the aphotic bathypelagic and abyssopelagic zones (>1000m, n = 17; Supplementary Table S1). Depth information was gathered from FishBase (Froese and Pauly, 2022) for all species based on documented observations. We binned diet data categorically as large evasive prey (e.g., fishes and squid), gelatinous zooplankton (e.g., jellyfishes, salps), or smaller zooplankton, following Friedman et al. (2019), with additional data gathered from FishBase (Froese and Pauly, 2022; Supplementary Table S1). We also assigned each species a tooth type category, adapted from Mihalitsis and Bellwood (2019). We defined four categories: edentulate (no visible teeth); microdont (single row of small, heterogenous, closely spaced teeth); macrodont (single row of relatively large, widely spaced teeth or combination of small and large teeth); and villiform (abundant, closely spaced bristle- or needle-like teeth in several rows or a single comb-like row). Feeding mode is poorly understood in many oceanic taxa due to the difficulty of observation and is generally inferred from morphology or phylogenetic bracketing, so we do not consider it in our analysis. We evaluated the correlation strength between shape and ecological and biomechanical data using distance-based regressions or permutational MANOVAs. This approach has been shown to be appropriate for high-dimensional morphometric data and was implemented with the function ‘procD.pgls’ (to account for phylogenetic non-independence) in the R package *geomorph* (Adams et al., 2023).

### Mechanical performance data

We calculated mechanical advantage (MA) for the lower jaw using the method outlined in Westneat (2004), where MA = L_in_/L_out_. Measurements were taken directly from the digital meshes for each specimen. The in-lever (L_in_) was measured as the distance from the centre of rotation of the quadrato-mandibular joint to the attachment of the adductor mandibularis on the coronoid process for jaw closing (MA_close_), and the attachment of the interopercular mandibular ligament on the retroarticular process for jaw opening MA_openA_ (Supplementary Fig. S3; Albertson et al., 2005). Due to the variable presence/absence of teeth across Pelagiaria, the out-lever (L_out_) was measured as the distance from the jaw joint to the anterior tip of the jaw rather than the tip of the distalmost tooth. This simplified MA model ignores the insertion angle of the adductor muscle and thus tends to overestimate force and underestimate velocity transmission but provides a useful comparative functional variable (Westneat, 2004). The four-bar linkage system that operates in the teleost oral jaw operates in the mandible between the quadrato-mandibular joint and the coronoid process via the coronoid-maxilla joint. Thus, when the lower jaw is depressed, force output is transmitted via this joint. For this reason, we also calculated a second opening MA value (MA_openB_) for each specimen to account for this connection. We took the in-lever for MA_openB_ as the same value for MA_openA_ (L_in_ opening), and measured the out-lever as the distance between the quadrato-mandibular joint and the attachment point of the ligament at the tip of the coronoid process (Motta, 1984).

We identified mandible morphotypes by assessing the clustering of specimens in morphospace using the ‘silhouette’ method, implemented using the ‘fviz_nbclust’ method in the R package *factoextra* (Kassambara and Mundt, 2020), using the first 35 PC scores (accounting for 95% of total shape disparity). We assessed differences in MA values between each morphotype with an ANOVA, and then evaluated these results using the ‘TukeyHSD’ function in the *stats* R package (R Core Team). This calculates the differences in observed means, with p-values adjusted for multiple tests.

We tested morphotype group separation within the morphospace using a discriminant function analysis (DFA) implemented with the ‘mvgls.dfa’ function in the R package *mvMORPH*, which assesses group distribution from a phylogenetic regression optimised for multivariate data (Clavel et al., 2015). This method also provides prediction power of the model and identifies the proportion of misassigned species per group.

### Evolutionary modelling

We used BayesTraitsV3 (Meade and Pagel, 2014) to determine the best-supported evolutionary model of the mandible, using phylogenetic PC (pPC) scores that accounted for >95% of combined total shape variation. We tested 10 alternative models: fixed and variable rate Brownian motion (BM) and Ornstein-Uhlenbeck (OU), with delta (**δ**), kappa (**κ**), and lambda (**λ**) tree transformations (Meade and Pagel, 2014). We determined the best-supported evolutionary model with Bayes Factor in the R package BTprocessR (Ferguson-Gow, 2021). We calculated mandible disparity through time (DTT) as a measure of relative subclade disparity using the ‘dtt’ function in the R package *geiger* v. 2.0.11 (Pennell et al., 2014), following Harmon et al. (2003). At each node on the tree, relative disparity is calculated as the mean of the relative disparities of all subclades whose ancestral lineages are present at that point in time by the disparity of the whole tree. Relative disparity values closer to zero indicate that morphological variation is mainly partitioned between subclades rather than within them. The empirical DTT curve, which we calculated from shape data, was compared to a null model of 2000 simulations of Brownian evolution calculated from tree topology (Harmon et al., 2003).

## Results

### Shape disparity and phylogenetic signal

The PCA performed on Procrustes-aligned mandible shape shows that PC 1 accounts for almost 50% of total shape disparity in the dataset. This axis of shape variation represents jaw elongation, with long, thin mandibles associated with negative PC values and short, deep mandibles associated with positive values (Fig. 1). PC 2 accounts for 16.8% of shape variation in the dataset, and is associated with robust, rounded mandible shape at positive values and an elongate triangular mandible shape at negative values. Chiasmodontidae, Scombridae and Gempylidae/Trichiuridae all occupy distinct regions of morphospace in Fig. 1, whereas Stromateidae, Centrolophidae, Bramidae and Caristiidae all show clear overlap along PCs 1 and 2. Phylogenetic signal in shape data is significant but low (K_mult_ = 0.34, *p* = 0.001), indicating a degree of shape convergence. Jaw shape is not significantly correlated with specimen centroid size after accounting for phylogeny (Z = 0.638, *p* = 0.264).

**Figure 1:**
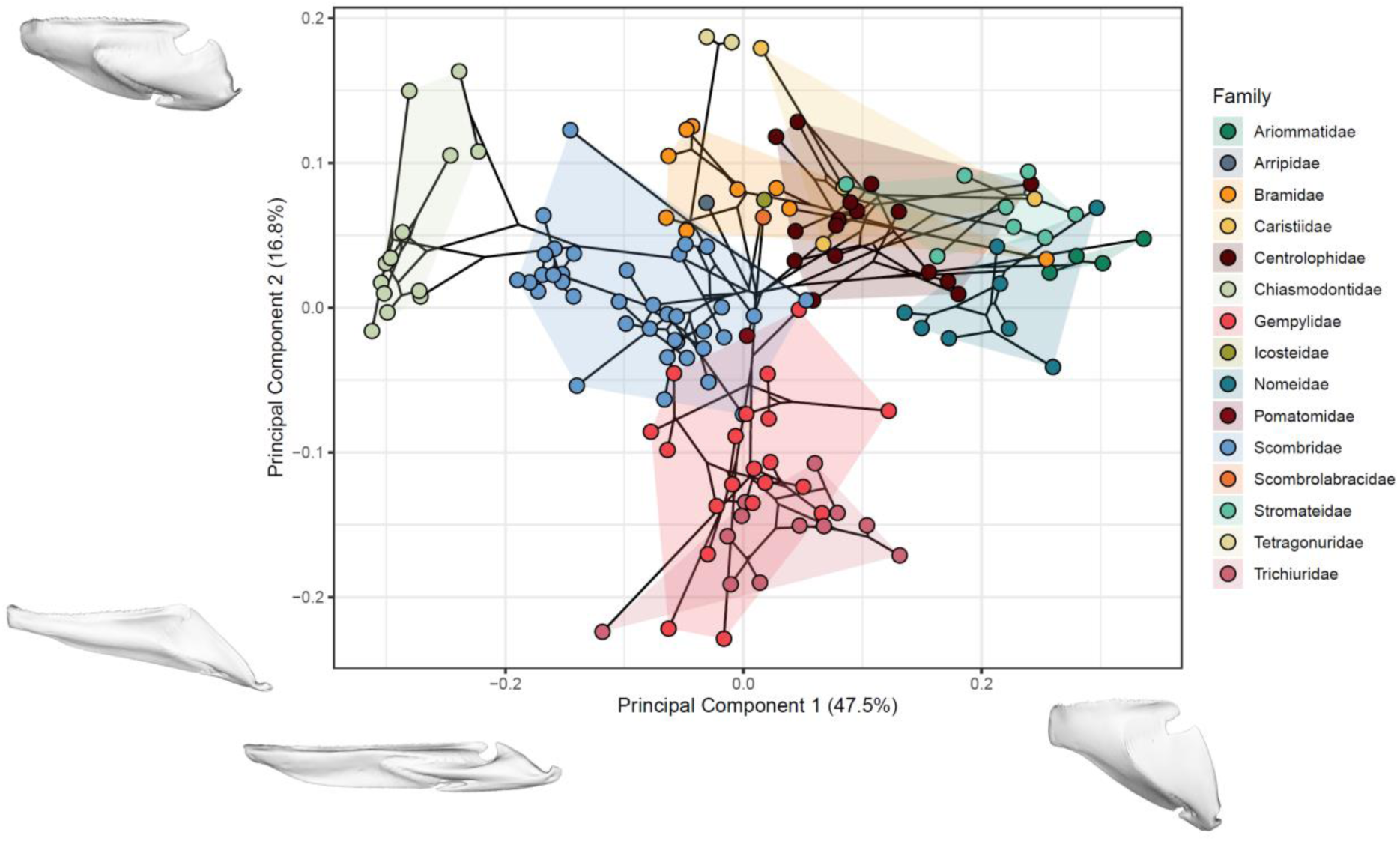
Phylomorphospace of pelagiarian mandibles. First two PCs are shown, with specimens coloured according to family. Convex hulls enclose members of each family within morphospace. Meshes at plot axes represent projected mandible shapes at maximum positive and negative values along the adjacent PCs.

Mandible shape is significantly correlated with habitat depth (Z = 2.58, *p* = 0.005; Fig. 2A). Shape is marginally correlated with tooth type (Z = 1.67, *p* = 0.048; Fig. 2D) and body elongation (Z = 1.65, *p* = 0.051; Fig. 2B) at the 5% significance level, but not with diet (Z = 0.90, *p* = 0.18; Fig. 2C; Supplementary Table S2).

**Figure 2:**
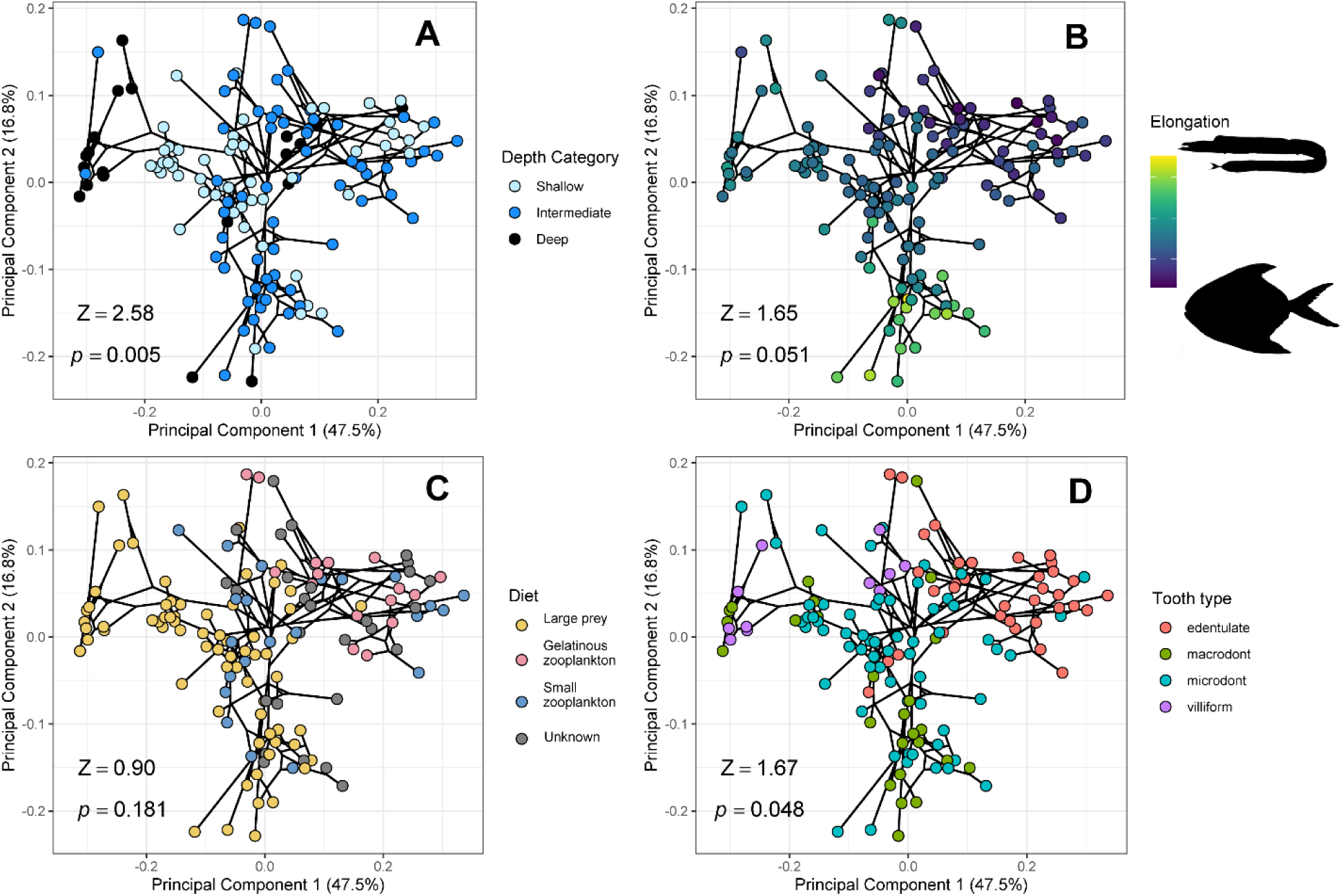
Phylomorphospaces of PCs 1 and 2 showing traits of pelagiarian mandibles. Shown are **A:** depth category; **B:** body elongation ratio**; C:** diet; **D**: tooth type. Effect size (Z) and p-value (p) are shown in each plot for phylogenetically corrected MANOVAs.

Mechanical advantage ranges from 0.039 (*Dysalotus alcocki*) to 0.245 (*Neocaristius heemstrai*) for MA_openA_, and from 0.151 (*Pseudoscopelus scutatus*) to 0.544 (*Ariomma regulus*) for MA_close_. Relative mechanical advantage of the pelagiarian mandible appears broadly consistent within families for both measures of jaw opening, and for jaw closing (Supplementary Fig. S5). MA_openB_ appears to show higher variation within and between families that other measures of MA. There are several discrepancies between relative levels of MA_openA_ and MA_close_ of individual species (Supplementary Fig. S6). Members of Trichiuridae and Gempylidae show low relative values for MA_openA_ but moderate to high values for MA_close_, suggesting that mandible shape in these families is optimised for high-velocity transmission when opening but strong force transmission when closing. Conversely, members of Caristiidae show relatively high MA_open_ but low MA_close_. Members of Scombridae and Chiasmodontidae show consistently low MA values for both opening and closing of the mandible, suggesting that jaw shape in these taxa has evolved for rapid movement.

Distribution of MA values in morphospace for jaw opening and closing are shown in Supplementary Fig. S7. Patterns of MA distribution appear similar for opening and closing, but there are notable differences between the plots. High MA_openA_ is largely restricted to the top right quadrant of the morphospace, with taxa in the central region and at negative values of PCs 1 and 2 showing correspondingly lower values (Supplementary Fig. S7A). Values of MA_openB_ follow similar patterns of distribution to those for MA_openA_, but with a slightly higher effect size (Z = 4.46, *p =* 0.001; Supplementary Fig. S7B). Conversely, MA_close_ shows an increase from the upper left quadrant to lower right quadrant (Supplementary Fig. S7C). Correlation of shape with MA is significant for each measure of MA, but effect size is highest for jaw closing (Z = 6.53, *p* = 0.001; Supplementary Table S2).

Six major morphospace clusters were revealed by the cluster analysis (Fig. 3). A discriminant function analysis of these groups revealed a very low training misclassification error rate of 0.7%, indicating good separation between clusters. MA values of each cluster are shown for MA_close_ (Fig. 3B) and MA_openA_ (Fig. 3C). Morphotypes 1 and 2 have significantly higher MA_close_ values than all other morphotypes, with 1 being the highest overall (Fig. 3B). Pairs of groups with similar MA_close_ values are formed by morphotypes 3 and 6, and 4 and 5, with 4 and 5 having the lowest MA_close_ values overall. Morphotype 1 also has the highest mean MA_openA_ value, but the MA_openA_ value for morphotype 3 is equally high (Fig. 3C). Contrastingly, the MA_openA_ value for morphotype 2 is low and is not significantly different to the values for groups 4, 5 and 6. Along with the significant correlation of MA with mandible shape, there is also a significant interaction between MA and mandible shape by morphotype (Z = 4.48, p = 0.001) for MA_close_, but not for MA_openA_ (Z = 1.29, p = 0.09).

**Figure 3:**
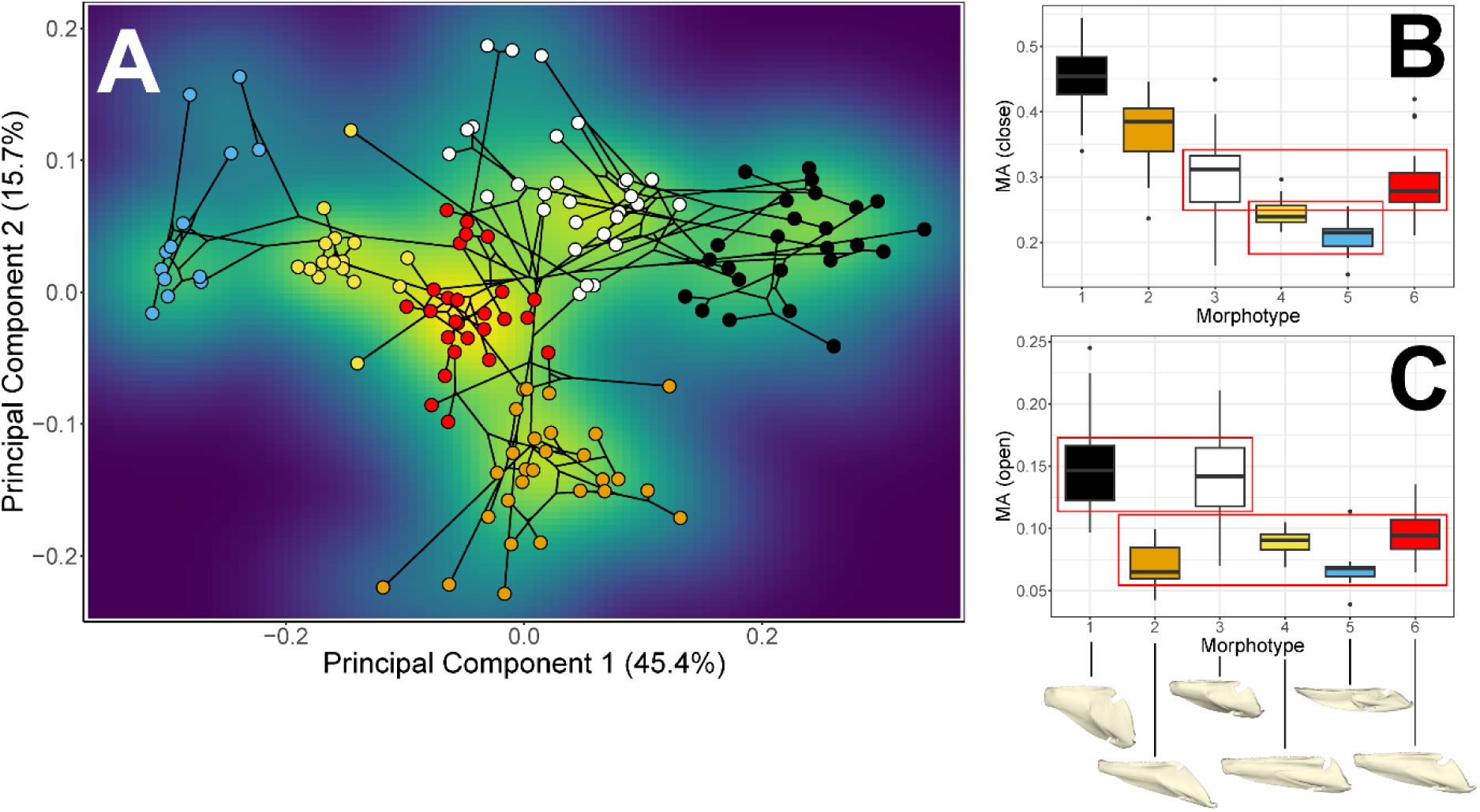
Mechanical advantage of major pelagiarian mandible morphotypes. Cluster analysis (**A**) of the six major mandible morphotypes. Mechanical advantage for each cluster is shown in bar plots for closing (**B**) and opening (**C**). Red boxes in **B** and **C** enclose morphotypes which do not significantly differ in MA. Mean shapes for each morphotype are shown below **C**.

We found strongest support for a single rate OU model of phenotypic evolution in the BayesTraits analysis (Supplementary Fig. S8). This mirrors the results of the same analysis in the pelagiarian neurocranium (Knapp et al., 2023), suggesting evolution towards a selective optimum but with little variation in overall rate.

Simulated DTT (Fig. 4, dashed line) shows an initial rapid increase in between-subclade disparity from the origin of the clade, with relative disparity accumulating gradually between subclades from the Eocene to the present day. In contrast, the observed data (Fig. 4, solid black line) shows more rapid accumulation of between-subclade disparity (steep negative slope falling outside the simulated model), indicating periods of rapid morphological divergence between clades and supporting adaptive radiation at the K/Pg boundary. Two further periods of between-subclade disparity accumulation occur in the late Pliocene and mid to late Pleistocene. These events are separated by prolonged periods of relatively little change in relative between-subclade disparity, especially between the late Eocene and early Pliocene, which fall well outside the simulated DTT.

**Figure 4:**
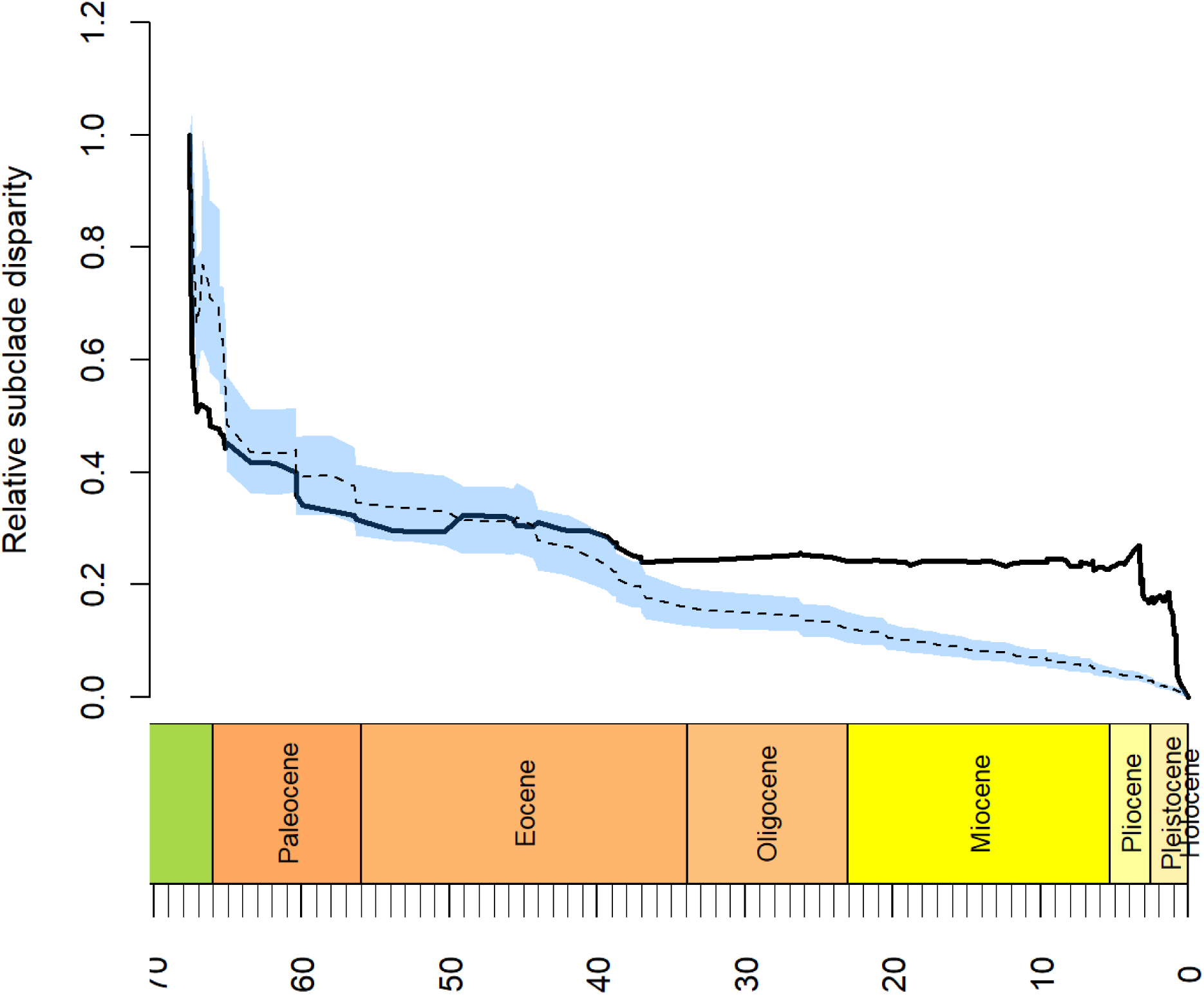
Pelagiarian mandible disparity through time plot. Observed average subclade disparity is shown by the solid black line. Negative slopes indicate that disparity is accumulating between, rather than within subclades. The median value of 2000 simulations of the null model of Brownian evolution is shown by the dashed black line, with the blue shaded area corresponding to 95% confidence intervals.

## Discussion

The pelagiarian mandible has rapidly evolved a high degree of both phenotypic and functional disparity. Our results support the hypothesis that the rapid diversification of families within Pelagiaria around the K/Pg mass extinction was driven by the vacating of ecological niches in the open ocean, and subsequent adaptive radiation. The strong association of mandible shape with mechanical advantage suggests that this disparity results from selection for mechanical performance in connection with the wide range of feeding strategies employed across Pelagiaria. Examination of MA within morphospace clusters (Fig. 3) reveals some similarities in mechanical performance in groups that have evolved very different feeding strategies and morphologies. Our results further emphasise the trophic complexity of the open ocean and demonstrate that this vast habitat can drive profound adaptive radiations in a way comparable with more structurally complex marine habitats such as coral reefs.

The low phylogenetic signal in mandible shape (K_mult_ = 0.34) is comparable with that of the pelagiarian neurocranium (K_mult_ = 0.27, Knapp et al., 2023). Low K_mult_ values suggest either a relatively low degree of shape diversification or a high amount of shape convergence/parallel evolution. This is consistent with an OU model, where shape is constrained along fewer axes of variation and mandible shape evolves within and between these optima, with convergent or parallel evolution resulting in low phylogenetic signal. The large amount of overlap between families in the upper right quadrant in Fig. 1 probably accounts for much of this, with taxa from Stromateidae, Centrolophidae, Bramidae and Caristiidae all occupying morphogroups 1 and 3 (Fig. 3). Methods exist for quantifying phenotypic convergence (Grossnickle et al., 2023; Stayton, 2015), but these are better suited to specific sets of a small number of taxa rather than large clades, which are extremely computationally expensive with current methods. It is worth noting that K_mult_ is measured across the entire dataset and that phenotypic convergence can vary considerably across such a large sample.

In contrast to the neurocranium (Knapp et al., 2023), the correlation of mandible shape with body elongation is weak in Pelagiaria. This suggests some relaxation of evolutionary integration between the neurocranium and mandible, with the neurocranium playing a key role in structuring the skull and in anchoring the suspensorium and axial muscles vital for powering suction feeding in teleosts (Camp et al., 2015), whereas the mandible has more freedom to explore shapes optimal for mechanical performance. This in turn highlights the important contribution of suspensorium shape in forming the fully functioning complex of elements which constitute the oral jaws, because it provides the fixed link in this system, connects it to the neurocranium and is an anchoring point for jaw musculature (Datavo and Vari, 2013; Westneat, 2004). The most elongate body shapes are found among Trichiuridae, but despite having relatively elongate jaws these taxa fall only midway along the axis that largely describes jaw elongation (Fig. 1). In contrast, the most elongate jaws in this dataset are found in Chiasmodontidae, which have evolved slender, widely gaping jaws like those of some other deep sea ambush predators (e.g., anglerfish; Heiple et al., 2023). The bodies of these taxa are only moderately elongated when compared with Trichiuridae, both in overall body shape and neurocranium shape (Knapp et al., 2023). Despite having important implications for locomotion mode and thus feeding strategy, the relationship between body shape and swimming performance is not linear (Friedman et al., 2021). For example, there is a trade-off between body length and body depth in minimising drag, and the optimum body shape for this is very close to the elongation ratio found in the Scombridae genera *Thunnus, Euthynnus, Katsuwonus* and *Auxis* (Altringham and Shadwick, 2001). Fast-swimming pelagiarian taxa are therefore likely to be found towards the middle of the body elongation continuum and tend to be a mixture of pursuit predators and ram-filter feeders (Fig. 2).

Diet is not significantly correlated with lower jaw shape in Pelagiaria. This may be surprising given the importance of the lower jaw in obtaining and processing food, but is likely explained by the relatively restricted dietary categories of this clade, combined with the different strategies employed to tackle similar prey across different families. For example, chiasmodontids and many scombrids feed on free-swimming, elusive prey but use entirely different strategies to do so. Chiasmodontids are deep sea ambush predators, whereas scombrids are generally fast-swimming pursuit predators in photic surface waters. Consequently, these groups have drastically diverged in almost every aspect of their biology. Many scombrids, however, have evolved to ram-feed on plankton in surface waters (e.g., mackerel, *Scomber*), where selection for efficient streamlining and agility to evade fast-swimming predators has maintained morphology very similar to predatory scombrids. Both predatory and planktivorous scombrids retain very similar mandible shapes (Fig. 2C), resulting from a lack of diversifying selection. The use of different feeding strategies at different depths likely explains the significant correlation between mandible shape and habitat depth, and feeding strategies are probably limited by the need to evade predators in surface waters.

In contrast to diet, the significant correlation of tooth type with mandible shape highlights the importance of tooth form in feeding strategy and suggests strong coevolution of these elements. For example, the large, barbed fangs of Trichiuridae (morphogroup 2; Bemis et al., 2018) work in conjunction with strong jaw musculature to seize, hold and break up prey (de Schepper et al., 2008). Taxa which filter feed (e.g., *Scomber*) or use ram-suction feeding (e.g., *Thunnus*) do not use their teeth for feeding in this way, and consequently, teeth are generally much reduced in size or absent entirely in these taxa (Fig. 3D). The teeth of the deep sea chiasmodontid taxa are often slender, sharp hooks, ideally suited for gripping prey while swallowing it whole.

Mechanical advantage is significantly correlated with jaw shape, with MA_close_ (i.e. bite force transmission) having an especially strong effect size. Mandible MA is well studied in teleost fishes, and although a basic measure of jaw performance, its implications for ecology have been widely discussed (Albertson et al., 2005; Wainwright and Richardson, 1995; Wainwright et al., 2004; Westneat, 1994 and 2004;). MA measurements from previous studies on different clades span a wide range of values (Table 1), reflecting trophic diversity in study taxa. The range of MA values in Pelagiaria is moderately high and comparable with studies investigating broad multi-clade samples across Caribbean reef fishes (Wainwright and Richard, 1995) and actinopterygians as a whole (Westneat, 2004) and has an upper limit comparable with that of the piranha *Serrasalmus rhombeus* (MA_close_ = 0.5; Grubich et al., 2012). This demonstrates the functional diversity of feeding mechanics in Pelagiaria. Other groups, e.g., Labridae (Wainwright et al., 2004, Table 1) show considerably higher MA_close_ values and ranges, driven by the high bite forces required in the highly specialised durophagous members of this clade.

**Table 1:** Summary of studies measuring mechanical advantage (MA) in fishes.

| Reference | Clade | n taxa | $MA_{openA}$ | $MA_{openA}$ | $MA_{openA}$ | $MA_{close}$ | $MA_{close}$ | $MA_{close}$ |
| --- | --- | --- | --- | --- | --- | --- | --- | --- |
|  |  |  | min | max | range | min | max | range |
| **This study** | **Pelagiaria** | **139** | **0.04** | **0.25** | **0.21** | **0.15** | **0.54** | **0.39** |
| Westneat 1994 | Labridae | 12 | 0.15 | 0.25 | 0.1 | 0.38 | 0.45 | 0.07 |
| Wainwright and Richard 1995 | Caribbean coral reef fishes | 34 | 0.08 | 0.32 | 0.24 | 0.1 | 0.5 | 0.4 |
| Westneat 2004 | Actinopterygii | 35 | 0.03 | 0.33 | 0.3 | 0.04 | 0.68 | 0.64 |
| Wainwright et al. 2004 | Labridae | 130 | 0.08 | 0.39 | 0.31 | 0.13 | 1.04 | 0.91 |

Combining MA values with shape and phylogenetic data reveals information that may not be apparent from measures of MA alone. MA_close_ is generally low across Scombridae and Chiasmodontidae, indicating selection for more rapid jaw closure but less powerful force transmission. This may be expected in both fast pursuit predators (Scombridae) and ambush predators (Chiasmodontidae), where rapid reactions are important in capturing elusive prey, despite their very different morphologies. MA_close_ is significantly higher in morphogroups 1 and 2 than in any other group (Fig. 3), despite being clearly separated from each other in morphospace. This indicates that high bite force transmission has evolved though very different shapes and feeding strategies. An analogous pattern of diverging morphology with convergent performance has been observed in fossorial mammals, indicating multiple solutions to the same adaptive challenge (Sansalone et al., 2020). Morphogroup 1 has the highest MA_close_ overall, with shape being characterised by short, deep jaws which are generally edentulate. Morphogroup 2 has lower MA_close_ than morphogroup 1, and its shape is characterised by long, triangular, toothed jaws. This group has thus evolved a unique jaw shape which combines elongation with high force transmission. Diet is also different between these two groups, with morphogroup 1 mainly feeding on gelatinous zooplankton and morphogroup 2 being mainly predators. Although diet is not significantly correlated with mandible shape across Pelagiaria, strong bite force transmission in these two groups may have evolved to deal with their specific prey. The gelatinous zooplankton dietary category is made up of large, soft-bodied animals, including hydrozoans, medusozoan and salps, which cannot often be swallowed whole. Fishes which feed on them must therefore bite chunks from their bodies or, in the case of some driftfish (Nomeidae), the tentacles and gonads of hydrozoans. Morphogroup 2 is entirely made up of taxa from Gempylidae and Trichiuridae, clades which possess large, sometimes barbed fangs with non-protrusible oral jaws, and are optimised for biting prey capture (Bemis et al., 2018; De Schepper et al., 2008). High bite force transmission has likely evolved in both groups because they generally feed on large prey which must be broken up before swallowing, contrasting with Chiasmodontidae (morphogroup 5), which swallow large prey whole and have the lowest MA_close_ of any group (Supplementary Fig. S5).

MA_open_ also varies significantly between morphogroups, but in a different way to MA_close_ (Fig. 3). Jaw opening is a more complicated process than jaw closing in teleosts and involves the kinetic interaction of more cranial elements (Albertson et al., 2005; Westneat, 2004). Morphogroups 2, 4, 5 and 6 have low MA_open_ values overall, and are not significantly different from one another. Morphogroups 1 and 3 have similar MA_open_ values, significantly higher than other groups. Differences in MA_open_ are not as straightforward to interpret as those for MA_close_, because the mandible does not contact with prey during opening and so it is not obvious why high MA (and hence force transmission) would be selected for. It is possible that selection for MA_close_ is translated to MA_open_ because of the limitations of jaw shape causing autocorrelation of inlever and outlever, but this is not true for all taxa in our study. For example, morphogroup 2 has significantly higher MA_close_ than all other morphogroups except morphogroup 1, but equally low MA_openA_ to all other morphogroups except 1 and 3. In the case of morphogroup 2, which is made up of members of Trichiuridae and Gempylidae, jaw shape appears to have been selected for high MA_close_ but low MA_openA_, allowing rapid jaw opening and forceful biting while maintaining an elongated shape.

The opening and closing mechanisms of the cichlid mandible are genetically modular and therefore able to evolve independently (Albertson et al., 2005), further implying that MA_open_ and MA_close_ are not always constrained by one another. Low MA_open_ enables high velocity transmission, which allows rapid response when capturing prey. It seems likely that, in the absence of any need for high force transmission when opening the jaw, higher MA_open_ would be selected against because it increases reaction time. The presence of taxa with significantly higher MA_open_ values suggests that high force transmission plays an important role in these taxa. This may relate to suction feeding, where high force transmission could be used to generate stronger pressure differentials needed to draw food into the mouth. There is, however, some disagreement among previous studies as to whether high MA_open_ (for generating rapid pressure change, e.g., Collar et al., 2022; Skorczewski et al., 2012) or low MA_open_ (for generating forceful pressure change, e.g., Albertson et al., 2005) is optimal for suction feeding.

No correlation between MA_open_ and jaw velocity was found in serranid fishes (Oufiero et al., 2012) and this could also be the case in Pelagiaria, though the former study used *in vivo* observations. Suction feeding in teleosts involves the interaction of many kinetic cranial and hyoid elements, and so a better understanding of the phenotypic coevolution of these other elements in Pelagiaria will be gained by incorporating them into future studies (Gartner et al., 2023). The transmission of force from the mandible to the upper jaw in the four-bar linkage mechanism of the teleost oral jaws (Motta, 1984) may also select for higher MA_openB_ in the mandible, and many pelagiarian taxa known to have non-protrusible upper jaws (e.g., Scombridae, Gempylidae) are found in morphogroups with very low MA_openB_, negating the requirement for strong force transmission when opening the jaws in these taxa. The significant correlation of mandible shape with MA_openB_ may add support for this, but presently little is understood about how force transmission via this route impacts suction feeding. Moreover, because of the nature of their open ocean habitat the feeding biology of many pelagiarian taxa is unknown, making assessments difficult for all but the best-known commercially important species.

We found that a single rate OU evolutionary model was best supported for the pelagiarian mandible, suggesting stabilising selection for optimum shape. Methods for testing for multiple optima with multivariate data are not well established at present, but the formation of clusters of taxa within morphospace (Fig. 3) suggests that the evolution of some clades (e.g., Chiasmodontidae, Gempylidae/Trichiuridae) that are associated with distinct clusters may have been driven by selection for shape optima. Simultaneously, clades which occupy multiple clusters or span wide regions of morphospace may be undergoing diversifying selection by evolving towards different shape optima. Some taxa, e.g., *Seriolella, Psenopsis, Peprilus, Cubiceps*, are members of families associated with a region of morphospace that shows high levels of overlap (Stromateidae, Centrolophidae, Bramidae and Caristiidae; Fig. 2), and indicates that some of these taxa may have undergone rapid adaptive evolution. Disparity through time (Fig. 4) shows rapid morphological diversification around the K/Pg boundary, adding support to the trophic diversification hypothesis of this clade and suggesting that mandible disparity in Pelagiaria was established early in the clade’s history. Although inferences of past disparity are hampered by the scarcity of fossil taxa from most of their evolutionary history, which may lead to an underestimate of past disparity and an overestimate of recent disparity change, this early rapid increase in disparity is still recoverable even with an extant-only dataset. Some very well-preserved pelagiarian fossils are available, particularly of scombrids, gempylids, and trichiurids, but their phylogenetic positions are currently not well resolved (Beckett and Friedman, 2016; Beckett et al., 2018; Monsch, 2004). Future work should integrate these examples with modern species to see if they amplify—or contradict—the pattern of rapid early divergence among lineages implied by extant taxa alone.

This study has shown that Pelagiaria has evolved a significant degree of disparity in both shape and mechanical performance of the mandible, and that this variation is likely to be associated with divergence in feeding strategy across different taxa. Reconstruction of clade-wide morphological disparity through time reveals periods of rapid evolutionary change in mandible shape, likely corresponding with adaptive radiations at the base of the clade and supporting the conclusions of Friedman et al. (2019). The ability of Pelagiaria to evolve such broad phenotypic diversity reveals that the open ocean is an environment well suited to driving rapid adaptive evolution, even across a relatively small clade with low dietary diversity. The teleost skull is complex both in terms of number of elements and in kinesis but although the mandible makes up only a part of these interconnected elements it plays a vital role in feeding, and even simple measures of mechanical performance have been shown to be broadly informative of function (Westneat 2005). Incorporating other kinetic elements of the suspensorium and oral jaws will allow us to examine the evolution of this clade in even greater detail, and to explore the effects of adaptive evolution on phenotypic integration and modularity across these complex structures.

## Acknowledgements

We are grateful to James Maclaine, Ollie Crimmen, and Emma Bernard (NHMUK), Peter Rask Möller (NHMD), Peter Bartsch and Edda Assel (MfN), Philippe Béarez and Jonathan Pfliger (MNHN), and Amanda Hay (AMS) for access to specimens. For assistance with scanning, we thank Vincent Fernandez and Brett Clark (NHMUK), Kristen Mahlow (MfN), Marta Bellato (MNHN), and Jenny Gibson (Royal National Orthopaedic Hospital NHS Trust). We also thank current and past members of the Goswami Lab at the Natural History Museum, London, for valuable feedback and discussion. This study includes data produced in the CTEES facility at University of Michigan, supported by the Department of Earth and Environmental Sciences and College of Literature, Science, and the Arts, and from the Department of Earth Sciences, University of Oxford, Oxford, UK. This study was funded by a Leverhulme Trust grant, no. RPG-2019-113.

