## Supplementary information for "Divergent morphologies with convergent performance in the mandible of pelagiarian fishes"

Andrew Knapp[1,2]*, Gizeh Rangel-de Lazaro[2,3], Matt Friedman[4], Zerina Johanson[2] , Kory M Evans[5], Sam Giles[6], Hermione T Beckett[7], and Anjali Goswami[2]

1. Centre for Integrative Anatomy, University College London, UK

2. Department of Science, Natural History Museum, London, UK.

3. School of Oriental and African Studies, University of London, UK.

4. Department of Earth and Environmental Sciences, University of Michigan, Ann Arbor, USA.

5. Department of Biosciences, Rice University, Houston, Texas, USA.

6. Department of Geography, Earth and Environmental Sciences, University of Birmingham, UK.

7. Department of Biology, King's High School for Girls, Warwick, UK

**Table S1: Information for all specimens used in study.** Sources of phylogenetic placement for all taxa are included in 'Phylogeny' column. Basic phylogeny was taken directly from Knapp et al., 2023, and the 23 taxa added for this study are highlighted in bold in the 'species' column. Additional taxa were placed on the phylogeny either by substituting a congeneric taxon or by grafting to known sister taxon, with source listed in the 'Phylogeny' column.

| Species | Family | Specimen number | Depth group | Diet | Elongation ratio | Tooth type | $MA_{openA}$ | $MA_{openB}$ | $MA_{close}$ | Phylogeny |
|---|---|---|---|---|---|---|---|---|---|---|
| *Acanthocybium solandri* | Scombridae | FMNH 44400 | shallow | large prey | 6.65 | microdont | 0.10 | 0.36 | 0.28 | Friedman et al. 2019 |
| *Allothunnus fallai* | Scombridae | NHMUK 2019.5.24.13 | shallow | large prey | 4.59 | edentulate | 0.10 | 0.33 | 0.29 | Miya et al. 2013 |
| *Aphanopus carbo* | Trichiuridae | NHMUK 2006.6.27.1 | deep | large prey | 16.11 | microdont | 0.06 | 0.26 | 0.24 | Friedman et al. 2019 |
| *Ariomma bondi* | Ariommatidae | UMMZ 228950 | intermediate | zooplankton | 3.55 | edentulate | 0.19 | 0.40 | 0.47 | Friedman et al. 2019 |
| *Ariomma indica* | Ariommatidae | UMMZ 219645 | intermediate | zooplankton | 2.45 | edentulate | 0.17 | 0.32 | 0.52 | Friedman et al. 2019 |
| *Ariomma melanum* | Ariommatidae | MCZ 168623 6 | intermediate | zooplankton | 4.15 | microdont | 0.15 | 0.29 | 0.50 | Friedman et al. 2019 |
| ***Ariomma regulus*** | Ariommatidae | NHMUK 1961.9.4.46.50.01 | intermediate | zooplankton | 2.26 | edentulate | 0.20 | 0.37 | 0.54 | Friedman et al. 2019 |
| *Arripis trutta* | Arripidae | UMMZ 209265 | shallow | large prey | 3.61 | villiform | 0.13 | 0.44 | 0.30 | Friedman et al. 2019 |
| *Auxis rochei* | Scombridae | UMMZ 250172 | shallow | zooplankton | 4.83 | edentulate | 0.08 | 0.26 | 0.30 | Friedman et al. 2019 |
| *Auxis thazard* | Scombridae | UMMZ 238740 | shallow | large prey | 4.44 | microdont | 0.09 | 0.35 | 0.26 | Friedman et al. 2019 |
| *Benthodesmus simyoni* | Trichiuridae | NHMUK 1972.1.10.64 | intermediate | NA | 33.68 | microdont | 0.05 | 0.13 | 0.38 | Friedman et al. 2019 |
| *Benthodesmus tenuis* | Trichiuridae | FMNH 88043 | intermediate | NA | 20.90 | microdont | 0.07 | 0.19 | 0.37 | Friedman et al. 2019 |
| *Brama brama* | Bramidae | FMNH 48931 | intermediate | large prey | 2.30 | macrodont | 0.14 | 0.53 | 0.26 | Arcila et al. 2021 |
| *Brama dussumieri* | Bramidae | NHMUK 2004.2.3.107 | shallow | NA | 2.10 | edentulate | 0.10 | 0.19 | 0.51 | Friedman et al. 2019 |
| *Brama orcini* | Bramidae | MfN 9949 | intermediate | NA | 1.95 | macrodont | 0.12 | 0.42 | 0.28 | Arcila et al. 2021 |
| ***Caristius fasciatus*** | Caristiidae | NHMD P40350 | intermediate | NA | 1.95 | microdont | 0.20 | 0.69 | 0.29 | Friedman et al. 2019 |
| *Caristius meridionalis* | Caristiidae | AMS 20071.038.31.002 | intermediate | NA | 1.87 | microdont | 0.20 | 1.23 | 0.16 | Arcila et al. 2021 |
| *Centrolophus niger* | Centrolophidae | NHMUK 2014.3.6.7 | deep | large prey | 3.91 | microdont | 0.15 | 0.46 | 0.32 | Miya et al. 2013 |
| *Chiasmodon niger* | Chiasmodontidae | UMMZ 228958 | deep | large prey | 7.33 | microdont | 0.06 | 0.29 | 0.22 | Friedman et al. 2019 |
| *Chiasmodon pluriradiatus* | Chiasmodontidae | NHMUK 1997.5.21.28 | deep | large prey | 6.15 | microdont | 0.07 | 0.34 | 0.22 | Friedman et al. 2019 |

| | | | | | | | | | | |
|---|---|---|---|---|---|---|---|---|---|---|
| *Cubiceps baxteri* | Nomeidae | NHMUK 1997.9.17.27 | shallow | jellyfish | 3.37 | microdont | 0.13 | 0.33 | 0.40 | Friedman et al. 2019 |
| *Cubiceps gracilis* | Nomeidae | NHMUK 2019.5.9.256 | shallow | jellyfish | 4.43 | edentulate | 0.12 | 0.26 | 0.46 | Arcila et al. 2021 |
| *Cubiceps pauciradiatus* | Nomeidae | NHMUK 2004.2.3.99.108 | intermediate | jellyfish | 3.92 | edentulate | 0.13 | 0.31 | 0.44 | Arcila et al. 2021 |
| *Cubiceps whiteleggi* | Nomeidae | NHMUK 1986.10.6:37-40 | intermediate | jellyfish | 3.37 | microdont | 0.12 | 0.30 | 0.41 | Friedman et al. 2019 |
| *Diplospinus multistriatus* | Gempylidae | YPM 0256719 | intermediate | zooplankton | 20.18 | macrodont | 0.07 | 0.17 | 0.40 | Friedman et al. 2019 |
| *Dysalotus alcocki* | Chiasmodontidae | NHMUK 2017.10.26.237 | deep | large prey | 5.9 | villiform | 0.04 | 0.16 | 0.25 | Friedman et al. 2019 |
| *Epinnula magistralus* | Gempylidae | UF 233577 | shallow | NA | 4.64 | macrodont | 0.10 | 0.25 | 0.39 | Friedman et al. 2019 |
| *Eumegistus illustris* | Bramidae | LACM 38573 | intermediate | zooplankton | 2.37 | villiform | 0.13 | 0.56 | 0.24 | Friedman et al. 2019 |
| *Eupleurogrammus glossodon* | Trichiuridae | NHMUK 1860.3.19.76 | shallow | large prey | 15.81 | macrodont | 0.04 | 0.11 | 0.40 | Beckett et al. 2018 |
| *Eupleurogrammus muticus* | Trichiuridae | NHMUK 1962_5_4_2 | shallow | large prey | 14.29 | macrodont | 0.05 | 0.14 | 0.39 | Beckett et al. 2018 |
| *Euthynnus affinis* | Scombridae | UMMZ 223134 | shallow | zooplankton | 4.03 | microdont | 0.09 | 0.35 | 0.27 | Friedman et al. 2019 |
| *Euthynnus alleteratus* | Scombridae | YPM M39708-71762 | shallow | large prey | 3.99 | microdont | 0.09 | 0.32 | 0.28 | Arcila et al. 2021 |
| ***Evoxymetopon taeniatus*** | Trichiuridae | UF 210626 | shallow | NA | 13.27 | macrodont | 0.06 | 0.13 | 0.45 | Friedman et al. 2019 |
| *Gasterochisma melampus* | Scombridae | NHMUK 1868.6.22.2 | intermediate | large prey | 3.69 | microdont | 0.12 | 0.43 | 0.27 | Friedman et al. 2019 |
| *Gempylus serpens* | Gempylidae | NHMUK 2019.5.9.257 | intermediate | large prey | 22.74 | microdont | 0.06 | 0.22 | 0.28 | Friedman et al. 2019 |
| *Grammatorcynus bicarinatus* | Scombridae | NHMUK 1872.4.6.25 | shallow | large prey | 5.2 | edentulate | 0.10 | 0.39 | 0.26 | Arcila et al. 2021 |
| *Grammatorcynus bilineatus* | Scombridae | MfN 8003 | shallow | large prey | 4.15 | microdont | 0.09 | 0.31 | 0.31 | Arcila et al. 2021 |
| *Gymnosarda unicolor* | Scombridae | FMNH 89962 | shallow | large prey | 3.89 | macrodont | 0.11 | 0.38 | 0.28 | Friedman et al. 2019 |
| *Hyperoglyphe japonica* | Centrolophidae | UMMZ 233106 | deep | zooplankton | 2.9 | edentulate | 0.18 | 0.39 | 0.45 | Friedman et al. 2019 |
| *Hyperoglyphe perciformis* | Centrolophidae | MNHN IC 1998-1430 | deep | large prey | 2.90 | edentulate | 0.17 | 0.60 | 0.28 | Friedman et al. 2019 |
| *Icichthys australis* | Centrolophidae | NHMUK 2009.5.28.12 | deep | NA | 2.63 | edentulate | 0.17 | 0.38 | 0.45 | Arcila et al. 2021 |
| ***Icosteus aenigmaticus*** | Icosteidae | NHMUK 1886.11.5.1 | intermediate | jellyfish | 5.14 | edentulate | 0.18 | 0.51 | 0.36 | Friedman et al. 2019 |
| *Kali indica* | Chiasmodontidae | NHMUK 2018.7.26.214_01 | intermediate | large prey | 2.78 | macrodont | 0.06 | 0.28 | 0.20 | Friedman et al. 2019 |
| *Kali kerberti* | Chiasmodontidae | FMNH 88149 | deep | large prey | 4.26 | macrodont | 0.06 | 0.24 | 0.25 | Friedman et al. 2019 |

| | | | | | | | | | | |
|---|---|---|---|---|---|---|---|---|---|---|
| *Kali macrodon* | Chiasmodontidae | NHMUK 1996.2.14.15.01 | deep | large prey | 8.06 | macrodont | 0.06 | 0.25 | 0.23 | Friedman et al. 2019 |
| *Katsuwonus pelamis* | Scombridae | FMNH 41899 | intermediate | zooplankton | 3.84 | edentulate | 0.08 | 0.32 | 0.26 | Friedman et al. 2019 |
| *Lepidocybium flavobrunneum* | Gempylidae | MNHN IC 2000.481 | deep | large prey | 4.54 | macrodont | 0.10 | 0.30 | 0.33 | Friedman et al. 2019 |
| *Lepidopus altifrons* | Trichiuridae | FMNH 64192 | intermediate | NA | 12.26 | macrodont | 0.05 | 0.12 | 0.43 | Friedman et al. 2019 |
| *Lepidopus caudatus* | Trichiuridae | NHMUK 1928.6.24.27 | intermediate | zooplankton | 16.57 | microdont | 0.05 | 0.14 | 0.40 | Friedman et al. 2019 |
| *Lepturacanthus savala* | Trichiuridae | UMMZ 219522 | shallow | large prey | 15.81 | macrodont | 0.06 | 0.21 | 0.30 | Friedman et al. 2019 |
| ***Neocaristius heemstrai*** | Caristiidae | AMS 25447 | intermediate | NA | 1.81 | edentulate | 0.24 | 0.57 | 0.43 | Friedman et al. 2019 |
| ***Neoepinnula americana*** | Gempylidae | FMNH 65717 | intermediate | NA | 4.32 | macrodont | 0.10 | 0.24 | 0.41 | Friedman et al. 2019 |
| *Neoepinnula orientalis* | Gempylidae | NHMUK 1986.9.8.164 | intermediate | large prey | 4.28 | macrodont | 0.10 | 0.24 | 0.39 | Friedman et al. 2019 |
| *Nesiarchus nasutus* | Gempylidae | FMNH 71422 | deep | large prey | 11.80 | macrodont | 0.07 | 0.22 | 0.31 | Friedman et al. 2019 |
| *Nomeus gronvii* | Nomeidae | NHMUK 1900.11.6.1 | intermediate | jellyfish | 3.63 | microdont | 0.18 | 0.39 | 0.46 | Froese and Pauly 2022 |
| *Orcynopsis unicolor* | Scombridae | NHMUK 1935.3.5.52 | shallow | large prey | 3.76 | microdont | 0.09 | 0.38 | 0.24 | Froese and Pauly 2022 |
| *Pampus argenteus* | Stromateidae | UMMZ 225567 | shallow | zooplankton | 1.45 | edentulate | 0.13 | 0.26 | 0.48 | Friedman et al. 2019 |
| ***Pampus chinensis*** | Stromateidae | NHMUK 1925.4.23.6.3 | shallow | jellyfish | 1.36 | edentulate | 0.16 | 0.43 | 0.36 | Friedman et al. 2019 |
| *Paradiplospinus antarcticus* | Gempylidae | LACM 10908 | deep | zooplankton | 12.89 | macrodont | 0.08 | 0.20 | 0.42 | Friedman et al. 2019 |
| *Paradiplospinus gracilis* | Gempylidae | NHMUK 2009.5.18.74 | intermediate | zooplankton | 9.70 | macrodont | 0.06 | 0.16 | 0.39 | Friedman et al. 2019 |
| *Peprilus alepidotus* | Stromateidae | UMMZ 199053 | shallow | jellyfish | 1.51 | edentulate | 0.12 | 0.24 | 0.51 | Friedman et al. 2019 |
| ***Peprilus burti*** | Stromateidae | UMMZ 249257 | shallow | jellyfish | 2.09 | edentulate | 0.12 | 0.26 | 0.46 | Arcila et al. 2021 |
| *Peprilus paru* | Stromateidae | UMMZ 145907 | shallow | jellyfish | 1.52 | edentulate | 0.16 | 0.69 | 0.24 | Friedman et al. 2019 |
| *Platyberyx opalescens* | Caristiidae | NHMUK 2005.4.28.1 | deep | NA | 2.05 | microdont | 0.21 | 0.88 | 0.24 | Arcila et al. 2021 |
| *Pomatomus saltatrix* | Pomatomidae | UMMZ 111069 | shallow | large prey | 3.81 | macrodont | 0.14 | 0.43 | 0.31 | Friedman et al. 2019 |
| *Promethichthys prometheus* | Gempylidae | UMMZ 250143 | intermediate | large prey | 7.67 | macrodont | 0.07 | 0.19 | 0.38 | Friedman et al. 2019 |
| ***Psenes arafurensis*** | Nomeidae | NHMUK 1997.9.17.10.01 | intermediate | NA | 2.55 | microdont | 0.15 | 0.44 | 0.34 | Friedman et al. 2019 |
| *Psenes cyanophrys* | Nomeidae | UMMZ 86219 | intermediate | NA | 2.10 | edentulate | 0.16 | 0.36 | 0.44 | Friedman et al. 2019 |
| *Psenes pellucidus* | Nomeidae | NHMUK 1963.5.14.481 | intermediate | zooplankton | 2.29 | edentulate | 0.12 | 0.24 | 0.51 | Arcila et al. 2021 |
| *Psenopsis anomala* | Centrolophidae | UMMZ 182977 | intermediate | zooplankton | 2.07 | microdont | 0.15 | 0.38 | 0.40 | Friedman et al. 2019 |
| *Psenopsis cyanea* | Centrolophidae | NHMUK 1937.6.28.1 | intermediate | NA | 2.93 | edentulate | 0.12 | 0.36 | 0.32 | Miya et al. 2013 |

| | | | | | | | | | | |
|---|---|---|---|---|---|---|---|---|---|---|
| *Psenopsis humerosa* | Centrolophidae | AUS I.24816-001 | intermediate | NA | 2.09 | edentulate | 0.16 | 0.36 | 0.44 | Froese and Pauly 2022 |
| *Psenopsis obscura* | Centrolophidae | NHMUK 2002.8.15.5 | intermediate | NA | 3.03 | edentulate | 0.15 | 0.35 | 0.43 | Froese and Pauly 2022 |
| *Pseudoscopelus altipinnis* | Chiasmodontidae | NHMUK 2004.8.17.39 | deep | large prey | 5.45 | villiform | 0.07 | 0.41 | 0.18 | Friedman et al. 2019 |
| *Pseudoscopelus astronesthidens* | Chiasmodontidae | MCZ 1614171 | deep | large prey | 6.45 | villiform | 0.07 | 0.31 | 0.22 | Friedman et al. 2019 |
| *Pseudoscopelus cordilluminatus* | Chiasmodontidae | NHMUK 2006.9.19.8 | deep | large prey | 4.05 | villiform | 0.07 | 0.33 | 0.21 | Melo 2010 |
| ***Pseudoscopelus obtusifrons*** | Chiasmodontidae | NHMUK 2004.8.17.26.02 | deep | large prey | 4.65 | villiform | 0.11 | 0.59 | 0.19 | Arcila et al. 2021 |
| *Pseudoscopelus pierbartus* | Chiasmodontidae | MCZ 164590 | deep | large prey | 4.62 | villiform | 0.07 | 0.38 | 0.18 | Friedman et al. 2019 |
| *Pseudoscopelus sagamianus* | Chiasmodontidae | NHMUK 1984.1.1.93 | intermediate | large prey | 5.89 | villiform | 0.07 | 0.31 | 0.22 | Melo 2010 |
| *Pseudoscopelus scutatus* | Chiasmodontidae | NHMUK 2004.8.17.24 | deep | large prey | 6.17 | villiform | 0.07 | 0.45 | 0.15 | Friedman et al. 2019 |
| *Pterycombus brama* | Bramidae | UF 168738 | intermediate | NA | 2.10 | microdont | 0.08 | 0.40 | 0.21 | Friedman et al. 2019 |
| *Pterycombus petersii* | Bramidae | MNHN 33644 | shallow | NA | 2.70 | microdont | 0.09 | 0.35 | 0.26 | Friedman et al. 2019 |
| *Rastrelliger faughni* | Scombridae | NHMUK 1984.1.18.208 | shallow | zooplankton | 4.00 | microdont | 0.07 | 0.28 | 0.25 | Arcila et al. 2021 |
| *Rastrelliger kanagurta* | Scombridae | UMMZ 226911 | shallow | zooplankton | 4.03 | microdont | 0.07 | 0.26 | 0.28 | Friedman et al. 2019 |
| ***Rexea alisae*** | Gempylidae | MNHN IC-1994-47 | intermediate | NA | 5.97 | macrodont | 0.06 | 0.14 | 0.44 | Friedman et al. 2019 |
| *Rexea antefurcata* | Gempylidae | NHMUK 1997.5.21.40 | intermediate | large prey | 6.04 | macrodont | 0.06 | 0.15 | 0.41 | Arcila et al. 2021 |
| *Rexea bengalensis* | Gempylidae | NHMUK 1996.9.25.36 | intermediate | large prey | 5.60 | macrodont | 0.09 | 0.21 | 0.43 | Arcila et al. 2021 |
| *Rexea prometheoides* | Gempylidae | FMNH 120779 | intermediate | large prey | 5.93 | macrodont | 0.09 | 0.22 | 0.38 | Friedman et al. 2019 |
| *Rexea solandri* | Gempylidae | UMMZ 216751 | intermediate | large prey | 5.09 | macrodont | 0.09 | 0.22 | 0.42 | Miya et al. 2013 |
| *Ruvettus pretiosus* | Gempylidae | NHMUK 1938.6.23 | intermediate | large prey | 5.04 | macrodont | 0.11 | 0.44 | 0.26 | Friedman et al. 2019 |
| *Sarda chilensis* | Scombridae | NHMUK 79.10.23.3 | shallow | large prey | 4.28 | macrodont | 0.09 | 0.34 | 0.27 | Arcila et al. 2021 |
| ***Sarda orientalis*** | Scombridae | NHMUK 1988.12.29.129.01 | shallow | large prey | 4.25 | macrodont | 0.08 | 0.36 | 0.22 | Arcila et al. 2021 |
| *Sarda sarda* | Scombridae | UMMZ 86218 | intermediate | NA | 2.33 | edentulate | 0.15 | 0.46 | 0.33 | Arcila et al. 2021 |
| ***Schedophilus maculatus*** | Scombridae | AMS 24645 | shallow | large prey | 3.99 | macrodont | 0.09 | 0.43 | 0.22 | Arcila et al. 2021 |
| ***Schedophilus ovalis*** | Centrolophidae | NHMUK 2011.2.25.1 | intermediate | jellyfish | 1.89 | microdont | 0.16 | 0.48 | 0.33 | Arcila et al. 2021 |
| ***Schedophilus pemarco*** | Centrolophidae | MNHN IC-1987-0982 | intermediate | NA | 2.41 | edentulate | 0.14 | 0.48 | 0.29 | Arcila et al. 2021 |

| | | | | | | | | | | |
|---|---|---|---|---|---|---|---|---|---|---|
| ***Schedophilus velaini*** | Centrolophidae | MNHN IC 2003-411 | intermediate | jellyfish | 3.01 | edentulate | 0.11 | 0.33 | 0.33 | Friedman et al. 2019 |
| *Scomber australasicus* | Scombridae | UMMZ 250152 | shallow | zooplankton | 5.32 | microdont | 0.08 | 0.28 | 0.30 | Friedman et al. 2019 |
| *Scomber japonicus* | Scombridae | UMMZ 250083 | shallow | zooplankton | 4.74 | microdont | 0.07 | 0.21 | 0.33 | Friedman et al. 2019 |
| *Scomber scombrus* | Scombridae | UMMZ 214543 | intermediate | zooplankton | 6.00 | microdont | 0.08 | 0.24 | 0.33 | Friedman et al. 2019 |
| ***Scomberomorus brasiliensis*** | Scombridae | NHMUK 1923.7.30.305.1 | shallow | large prey | 5.05 | macrodont | 0.09 | 0.40 | 0.23 | Arcila et al. 2021 |
| *Scomberomorus cavalla* | Scombridae | MfN 1445 | shallow | large prey | 4.19 | macrodont | 0.09 | 0.38 | 0.25 | Arcila et al. 2021 |
| ***Scomberomorus commerson*** | Scombridae | NHMD P7497 | shallow | large prey | 5.56 | macrodont | 0.09 | 0.39 | 0.24 | Miya et al. 2013 |
| *Scomberomorus guttatus* | Scombridae | MfN 1442 | shallow | large prey | 4.77 | macrodont | 0.10 | 0.38 | 0.26 | Arcila et al. 2021 |
| ***Scomberomorus koreanus*** | Scombridae | NHMUK 1983.12.13.28 | shallow | large prey | 4.37 | macrodont | 0.10 | 0.40 | 0.26 | Arcila et al. 2021 |
| *Scomberomorus lineolatus* | Scombridae | MNHN IC A 5802 | shallow | large prey | 4.25 | microdont | 0.09 | 0.39 | 0.22 | Jeena et al. 2022 |
| *Scomberomorus maculatus* | Scombridae | UMMZ 199143 | shallow | large prey | 4.53 | macrodont | 0.09 | 0.37 | 0.24 | Friedman et al. 2019 |
| *Scomberomorus niphonius* | Scombridae | UMMZ 167374 | shallow | large prey | 6.46 | macrodont | 0.07 | 0.29 | 0.24 | Friedman et al. 2019 |
| *Scomberomorus regalis* | Scombridae | UMMZ 143137 | shallow | large prey | 5.61 | macrodont | 0.08 | 0.31 | 0.26 | Friedman et al. 2019 |
| *Scomberomorus sierra* | Scombridae | ZMB 15569 | shallow | large prey | 5.49 | macrodont | 0.11 | 0.49 | 0.22 | Arcila et al. 2021 |
| ***Scomberomorus sinensis*** | Scombridae | MNHN IC A 5804 | shallow | large prey | 4.23 | macrodont | 0.07 | 0.32 | 0.22 | Arcila et al. 2021 |
| *Scomberomorus tritor* | Scombridae | MfN 9410 | shallow | large prey | 4.27 | macrodont | 0.09 | 0.37 | 0.23 | Arcila et al. 2021 |
| *Scombrolabrax heterolepis* | Scombrolabracidae | UF 167280 | intermediate | large prey | 4.50 | macrodont | 0.11 | 0.33 | 0.35 | Friedman et al. 2019 |
| *Seriolella brama* | Centrolophidae | NHMUK 1873.12.13.54 | intermediate | jellyfish | 2.83 | edentulate | 0.11 | 0.28 | 0.39 | Arcila et al. 2021 |
| *Seriolella caerulea* | Centrolophidae | NHMUK 2019.5.24.4 | shallow | jellyfish | 2.11 | edentulate | 0.15 | 0.43 | 0.36 | Arcila et al. 2021 |
| ***Seriolella porosa*** | Centrolophidae | NHMUK 1936.8.26.1062.6.1 | intermediate | NA | 3.60 | microdont | 0.12 | 0.32 | 0.38 | Arcila et al. 2021 |
| *Stromateus brasiliensis* | Stromateidae | NHMUK 2019.5.24.1 | shallow | NA | 2.23 | edentulate | 0.14 | 0.33 | 0.43 | Miya et al. 2013 |
| *Stromateus fiatola* | Stromateidae | NHMUK 1920.9.7.3 | shallow | large prey | 2.36 | edentulate | 0.11 | 0.28 | 0.41 | Friedman et al. 2019 |
| *Stromateus stellatus* | Stromateidae | NHMUK 1936.8.26.1072 | shallow | NA | 2.46 | edentulate | 0.22 | 0.49 | 0.46 | Miya et al. 2013 |
| *Taractes asper* | Bramidae | NHMUK 1953.10.28.2 | shallow | large prey | 2.62 | microdont | 0.12 | 0.46 | 0.27 | Friedman et al. 2019 |
| *Taractes rubescens* | Bramidae | MCZ 148060 | intermediate | zooplankton | 2.49 | macrodont | 0.14 | 0.61 | 0.23 | Friedman et al. 2019 |
| *Taractichthys* | Bramidae | FMNH 63863 | intermediate | NA | 1.62 | villiform | 0.15 | 0.58 | 0.25 | Friedman et al. 2019 |

| | | | | | | | | | | |
|---|---|---|---|---|---|---|---|---|---|---|
| *steindachneri* | | | | | | | | | | |
| *Tentoriceps cristatus* | Trichiuridae | NHMUK 1987.1.23.28 | shallow | large prey | 21.43 | macrodont | 0.06 | 0.16 | 0.40 | Friedman et al. 2019 |
| *Tetragonurus cuvieri* | Tetragonuridae | NHMD I 39502-001 | intermediate | jellyfish | 6.25 | microdont | 0.11 | 0.35 | 0.31 | Friedman et al. 2019 |
| *Tetragonurus sp* | Tetragonuridae | NHMUK 1998.8.9.9520 | intermediate | jellyfish | 6.86 | microdont | 0.09 | 0.41 | 0.23 | Friedman et al. 2019 |
| *Thunnus alalunga* | Scombridae | NHMUK 2004.11.1.299 | intermediate | large prey | 3.54 | edentulate | 0.11 | 0.35 | 0.30 | Arcila et al. 2021 |
| *Thunnus albacares* | Scombridae | NHMUK 1973.6.6.1 | shallow | large prey | 3.38 | microdont | 0.13 | 0.41 | 0.31 | Arcila et al. 2021 |
| *Thunnus atlanticus* | Scombridae | NHMUK 2012.8.15.13 | intermediate | large prey | 3.39 | microdont | 0.12 | 0.41 | 0.30 | Miya et al. 2013 |
| *Thunnus orientalis* | Scombridae | FMNH 58745 | intermediate | large prey | 3.36 | microdont | 0.10 | 0.35 | 0.28 | Friedman et al. 2019 |
| *Thunnus tonggol* | Scombridae | UMMZ 225078 | intermediate | large prey | 4.41 | microdont | 0.12 | 0.43 | 0.27 | Friedman et al. 2019 |
| *Thyrsites atun* | Gempylidae | NHMUK 1927.12.6.75 | intermediate | large prey | 7.94 | microdont | 0.08 | 0.21 | 0.37 | Arcila et al. 2021 |
| *Thyrsitoides marleyi* | Gempylidae | NHMUK 1986.9.8.147 | intermediate | large prey | 8.66 | macrodont | 0.08 | 0.28 | 0.31 | Friedman et al. 2019 |
| ***Thyrsitops lepidopoides*** | Gempylidae | NHMD P7353 | intermediate | large prey | 4.45 | macrodont | 0.08 | 0.26 | 0.33 | Arcila et al. 2021 |
| *Tongaichthys robustus* | Gempylidae | MNHN IC-2000-0460 | intermediate | NA | 4.28 | microdont | 0.10 | 0.25 | 0.40 | Beckett et al. 2018 |
| ***Trichiurus gangeticus*** | Trichiuridae | NHMD P73184 | intermediate | large prey | 13.00 | macrodont | 0.04 | 0.13 | 0.34 | Miya et al. 2021 |
| *Trichiurus lepturus* | Trichiuridae | UMMZ 219710 | intermediate | large prey | 13.86 | macrodont | 0.06 | 0.17 | 0.34 | Friedman et al. 2019 |
| *Tubbia tasmanica* | Centrolophidae | AMS IB.1148 | intermediate | NA | 3.16 | edentulate | 0.17 | 0.54 | 0.31 | Miya et al. 2013 |

**Abbreviations: AMS:** Australian Museum, Sydney; **FMNH:** Field Museum of Natural History; **LACM:** Los Angeles County Museum; **MCZ:** Museum of Comparative Zoology, Harvard University; **MfN:** Museum für Naturkunde, Berlin; **MNHN:** Muséum National d'Histoire Naturelle, Paris; **NHMD:** Natural History Museum of Denmark, Copenhagen; **NHMUK:** Natural History Museum, London; **UF:** Florida Museum of Natural History; **UMMZ:** University of Michigan Museum of Zoology; **YPM:** Yale Peabody Museum.

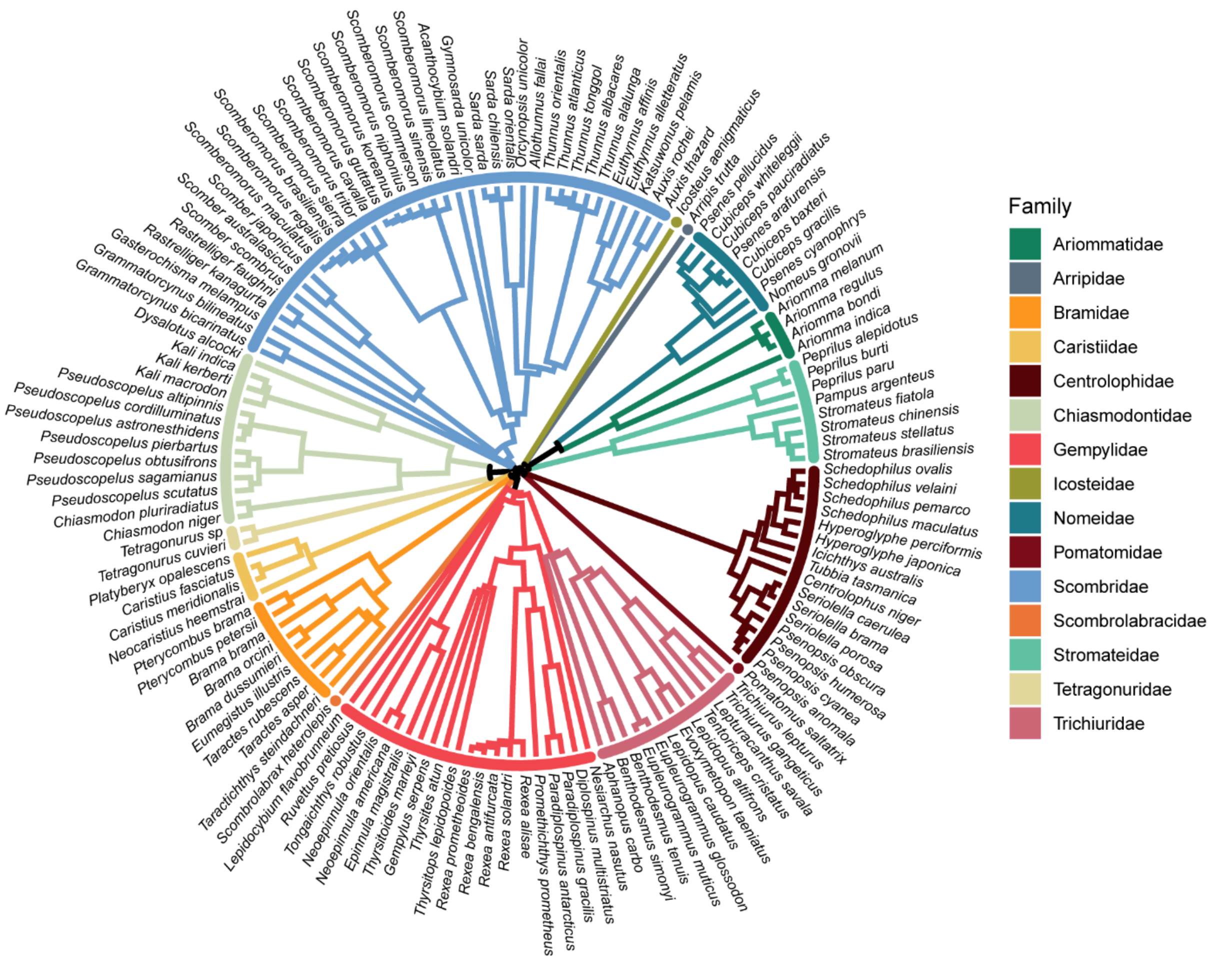

**Figure S1: Phylogeny of taxa used in this study.** Branches are coloured according to family.

### Mandible landmark list

### Anatomical landmarks

1. Dorsal point of symphysis on midline of dentary
2. Ventral point of symphysis on midline of dentary
3. Posterior tip of dentary at lowermost contact with articular
4. Posterior tip of toothrow of dentary
5. Anterior ventral process of articular
6. Midline of posterior margin of jaw joint on articular
7. Midline of anterior margin of jaw joint on articular
8. Tip of coronoid process

**Semilandmark curves**

1. Points 1-2 along dentary symphysis
2. Points 2-3 along ventral margin of dentary
3. Points 3-4 along labial dentary/articular suture
4. Points 4-1 along labial margin of tooth row
5. Points 4-1 along lingual margin of tooth row
6. Points 4-3 along lingual dentary/articular suture
7. Points 5-6 along ventral margin of articular
8. Points 6-7 along labial margin of jaw joint
9. Points 7-8 along dorsal margin of coronoid process of articular
10. Points 8-5 along lingual articular/dentary suture

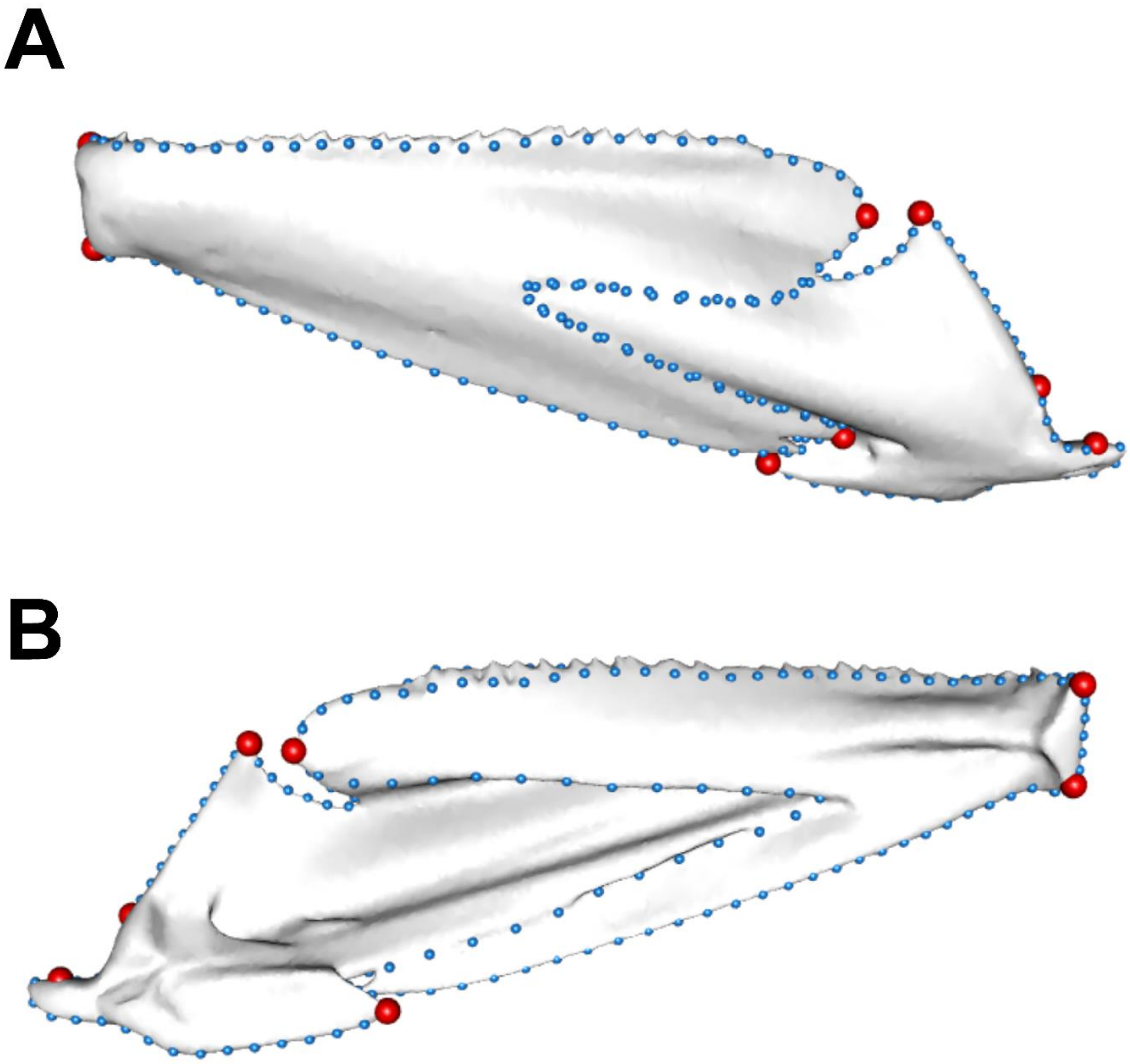

**Figure S2: Landmark guide for Pelagiaria mandible.** Landmarks are shown on the labial (A) and lingual (B) surfaces of the mandible, with anatomical landmarks in red and semilandmark curves in blue.

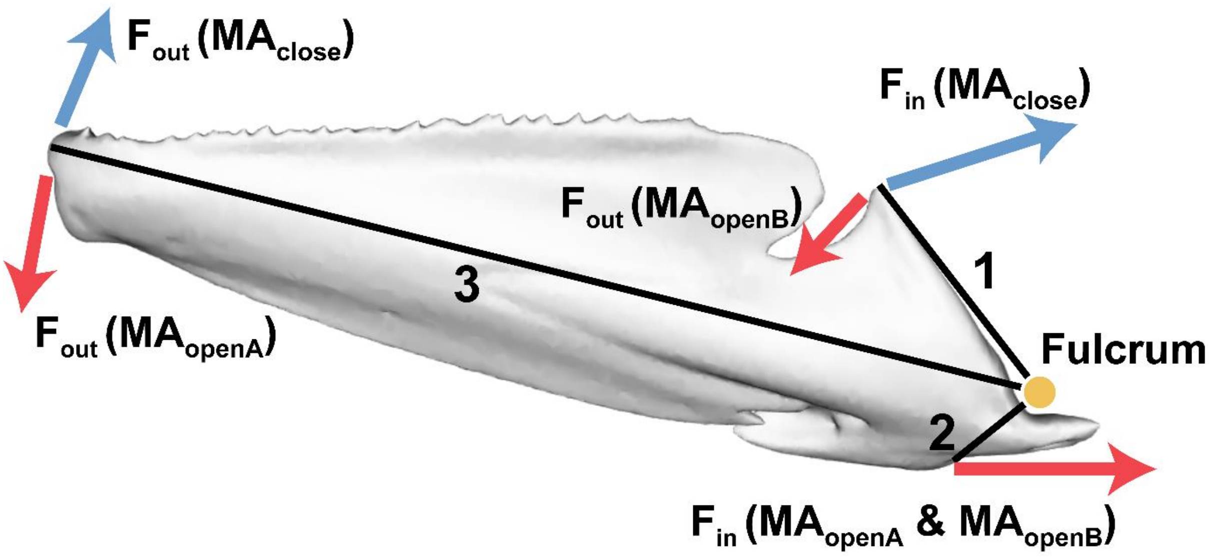

**Figure S3: Schematic of jaw lever measurements for calculating mechanical advantage (MA), show on mandible of *Thunnus tonggol*.** Solid black lines (**1**, **2**, **3**) represent lever arms: **1** = in-lever for jaw closing; **2** = in-lever for jaw opening; **3** = out-lever for jaw closing and jaw opening. Lever arm **1** also acts as the out lever for the calculation of $MA_{openB}$. Yellow circle represents the fulcrum (i.e. centre of rotation of quadrato-mandibular joint). Coloured arrows represent force inputs ($F_{in}$) and outputs ($F_{out}$), with red arrows representing jaw opening and blue arrows representing jaw closing.

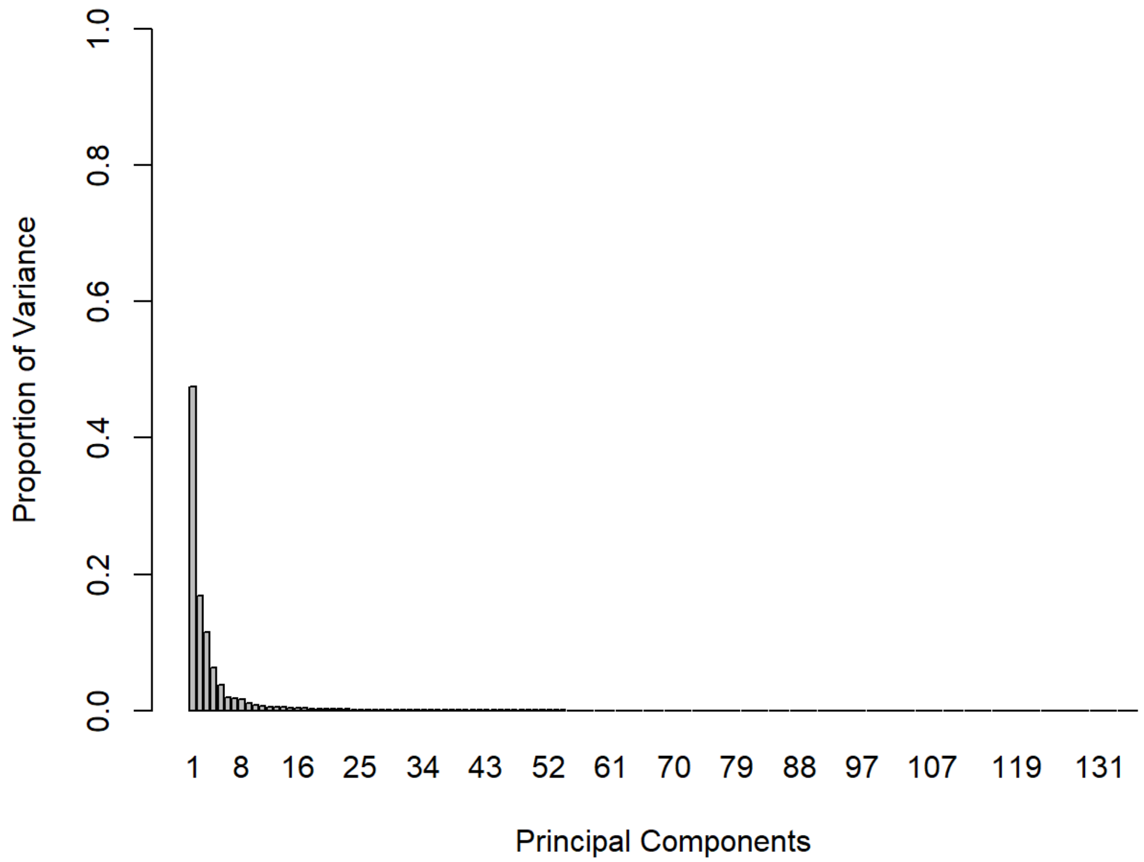

**Figure S4: Scree plot of PC contributions to shape variance.**

**Table S2: Results of allometry analysis, MANOVAs for categorical traits, and phylogenetic regressions for MA values.**

| | Rsq | F | Z | p |
|---|---|---|---|---|
| **Allometry** | 0.009 | 1.185 | 0.638 | 0.264 |
| **Depth** | 0.037 | 2.550 | 2.577 | 0.005 |
| **Elongation** | 0.016 | 2.257 | 1.653 | 0.051 |
| **Tooth type** | 0.035 | 1.606 | 1.672 | 0.048 |
| **Diet** | 0.025 | 1.326 | 0.898 | 0.181 |
| **$MA_{openA}$** | 0.032 | 4.506 | 3.326 | 0.003 |
| **$MA_{openB}$** | 0.059 | 8.452 | 3.732 | 0.001 |
| **$MA_{close}$** | 0.118 | 18.109 | 8.114 | 0.001 |

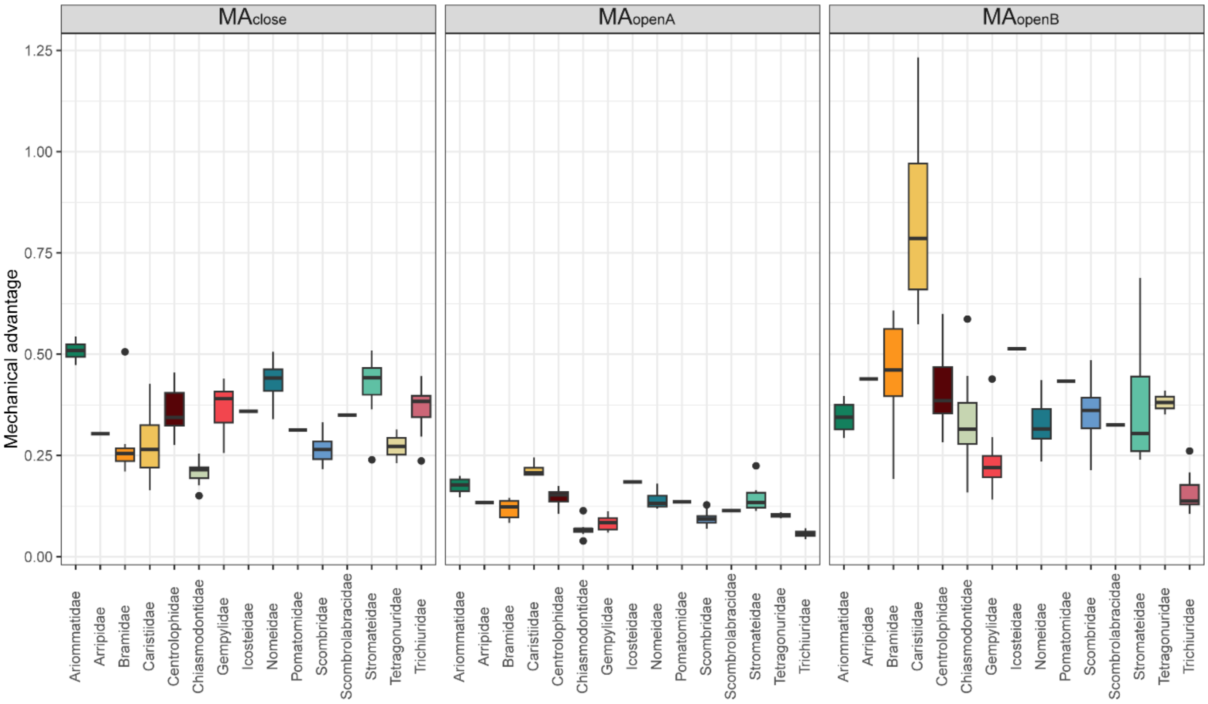

**Figure S5: Per-family MA values.** $MA_{close}$ (left plot), $MA_{openA}$ (middle plot) and $MA_{openB}$ (right plot) are shown for each Pelagiarian family in this study.

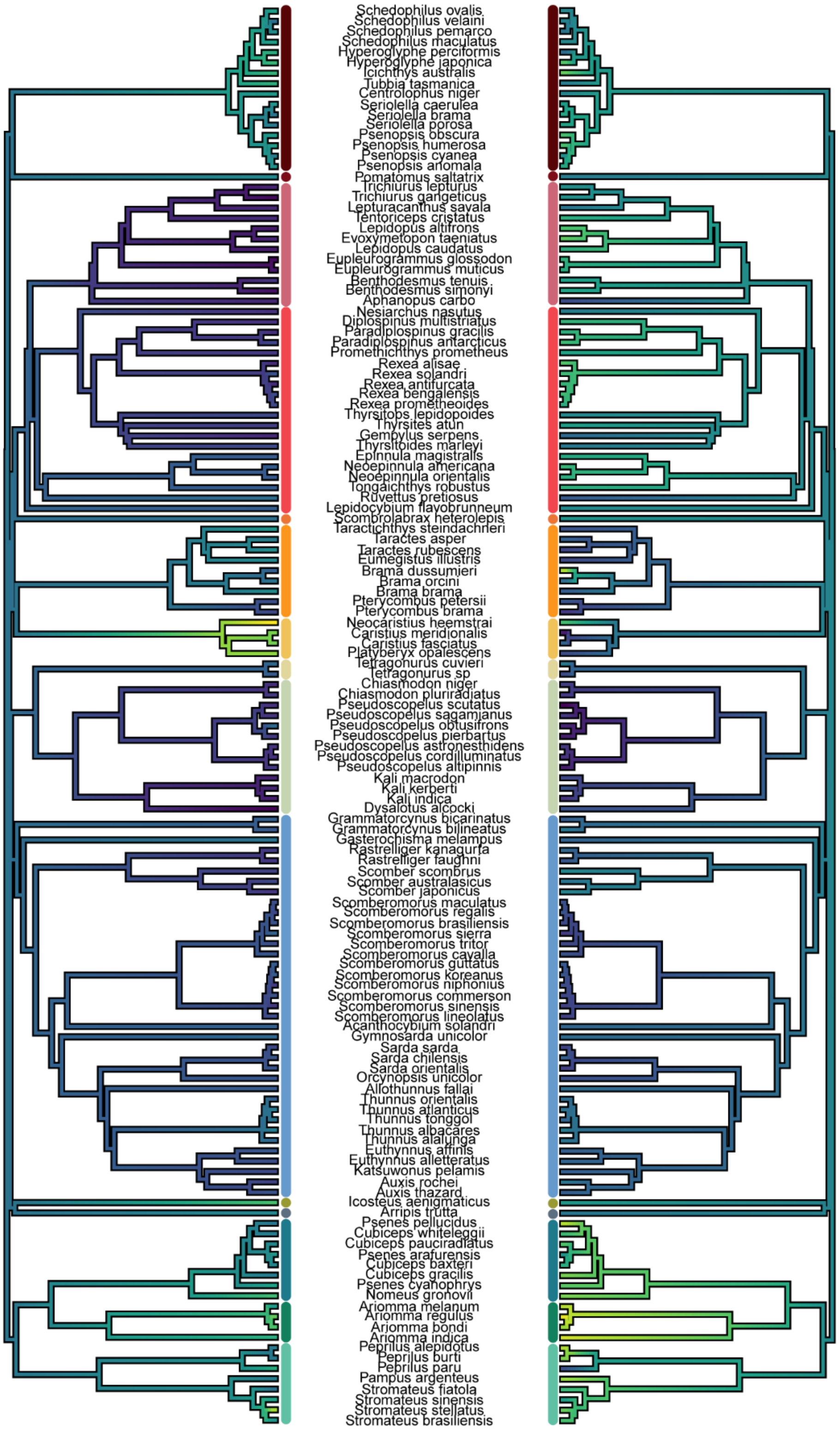

8

**Figure S6: Phylogeny of study sample with MA values.** MA values are shown for opening (left tree) and closing (right tree). Solid vertical bars at branch tips delimit families (see Supplementary Figure S1).

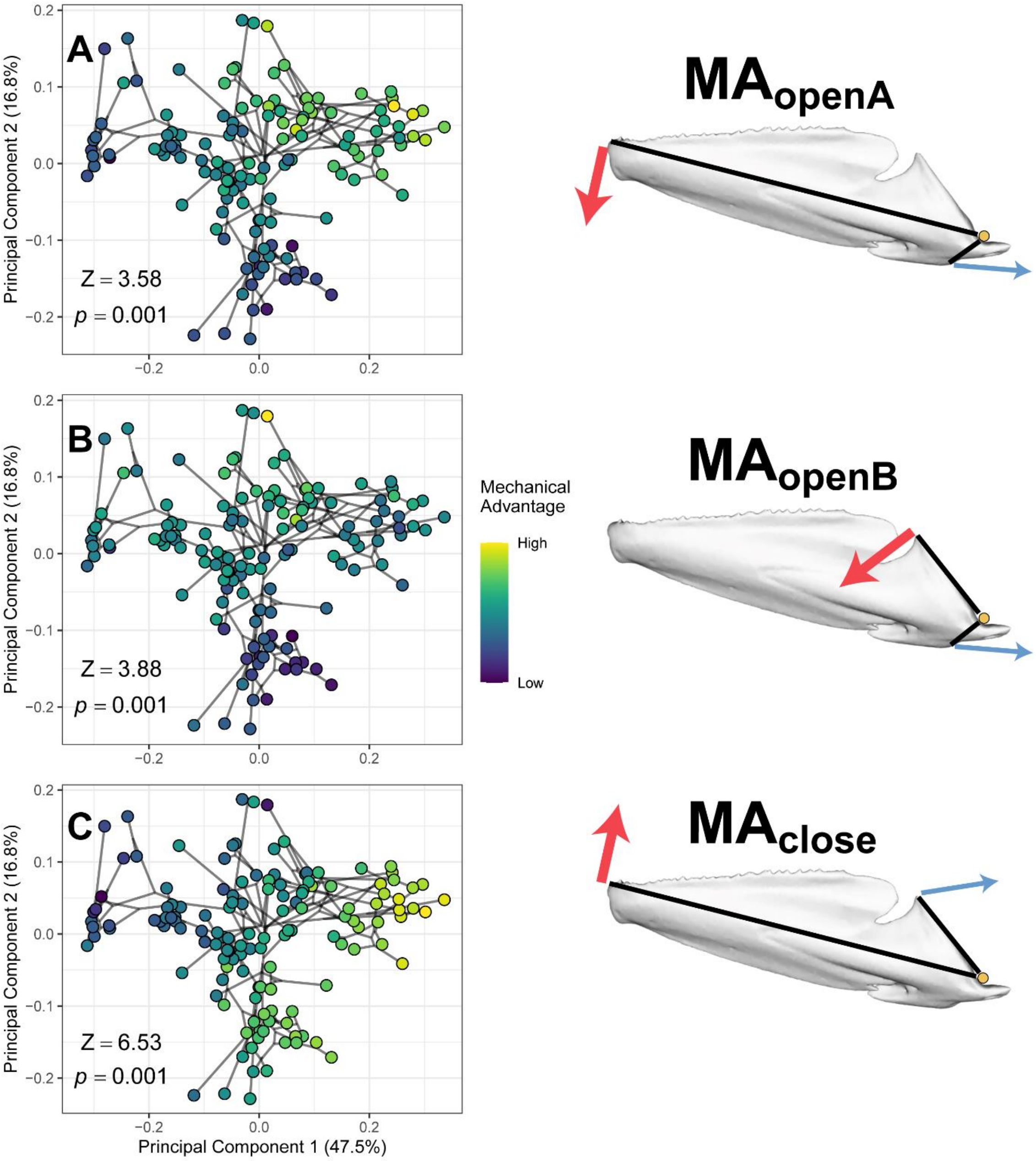

**Figure S7: Mechanical advantage (MA) measurements for pelagiarian jaw opening A (A), opening B (B) and closing (C).** Lighter colours represent higher MA values, darker colours represent lower MA values. Schematics show MA being measured for each plot. Blue arrows show force input; red arrows show force output; Solid black lines represent lever arms, and yellow circle represents fulcrum (i.e. centre of rotation of quadrato-mandibular joint. See Supplementary Fig. S1 for description of MA measurements).

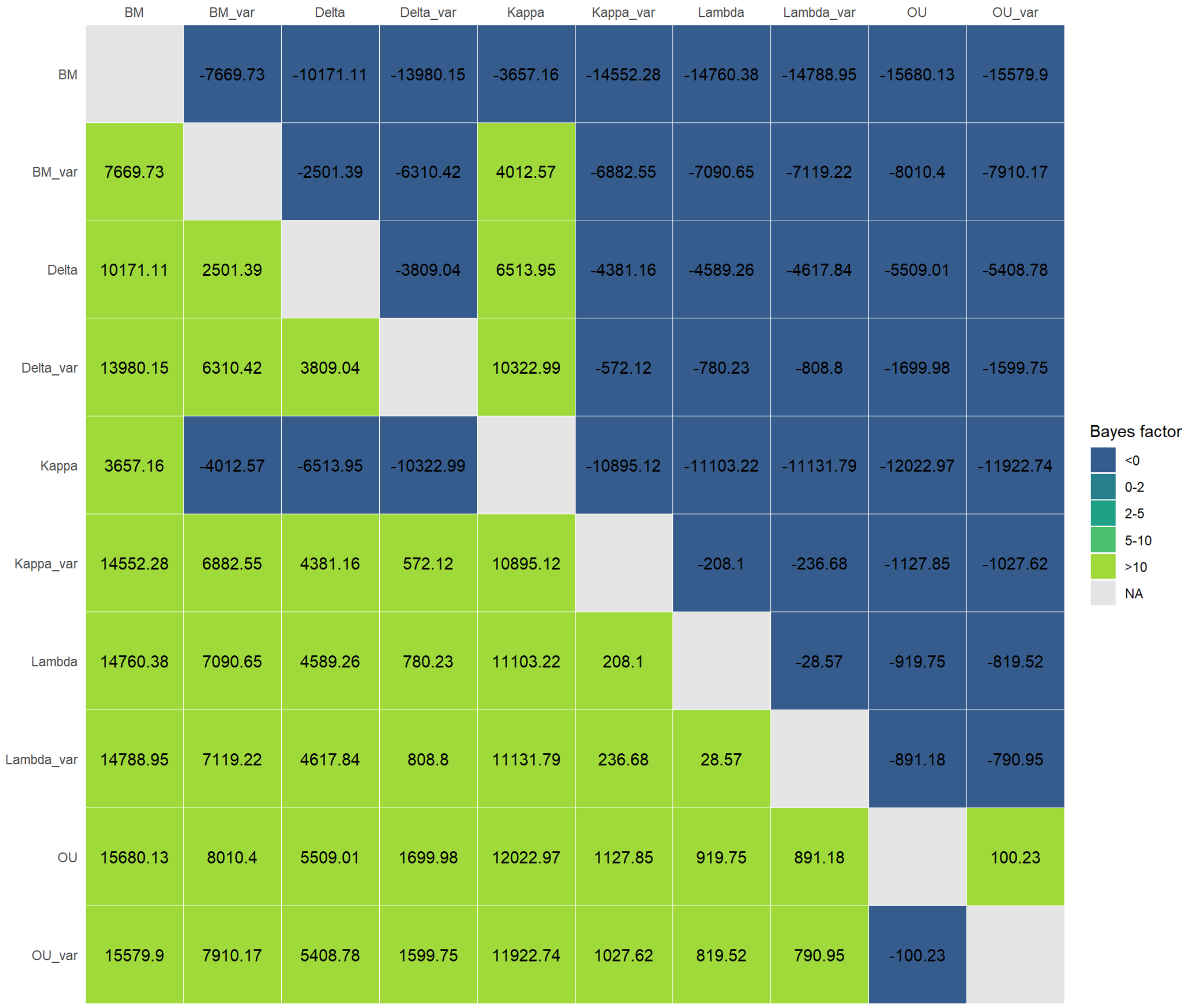

**Figure S8: Results of BayesTraits analysis.** Boxes contain pairwise marginal likelihood values. Higher values indicate better support. Each model is read horizontally, i.e. values in rows represent each model's marginal likelihood value with the corresponding model in each column.
